# The CDK8 inhibitor DCA promotes a tolerogenic chemical immunophenotype in CD4^+^ T cells via a novel CDK8-GATA3-FOXP3 pathway

**DOI:** 10.1101/855429

**Authors:** Azlann Arnett, Keagan G Moo, Kaitlin J Flynn, Thomas B Sundberg, Liv Johannessen, Alykhan F Shamji, Nathanael S Gray, Thomas Decker, Ye Zheng, Vivian H Gersuk, David E Levy, Isabelle J Marié, Ziaur S Rahman, Peter S Linsley, Ramnik J Xavier, Bernard Khor

## Abstract

Immune health requires innate and adaptive immune cells to engage precisely balanced pro- and anti-inflammatory forces. We employ the concept of chemical immunophenotypes to classify small molecules functionally or mechanistically according to their patterns of effects on primary innate and adaptive immune cells. The high-specificity, low-toxicity cyclin dependent kinase 8 (CDK8) inhibitor DCA exerts a distinct tolerogenic profile in both innate and adaptive immune cells. DCA promotes T_reg_ and Th2 differentiation, while inhibiting Th1 and Th17 differentiation, in both murine and human cells. This unique chemical immunophenotype led to mechanistic studies showing that DCA promotes T_reg_ differentiation in part by regulating a previously undescribed CDK8-GATA3-FOXP3 pathway that regulates early pathways of Foxp3 expression. These results highlight previously unappreciated links between T_reg_ and Th2 differentiation and extend our understanding of the transcription factors that regulate T_reg_ differentiation and their temporal sequencing. These findings have significant implications for future mechanistic and translational studies of CDK8 and CDK8 inhibitors.

## Introduction

The immune system comprises innate and adaptive immune cells whose collaborative and coordinated responses maintain the healthy state. Each cell type can exert either pro- or anti- inflammatory forces. For example, innate immune cells can secrete either pro- (e.g. IFNγ) or anti- (e.g. IL-10) inflammatory cytokines; similarly, CD4^+^ T cells can differentiate into either pro- (e.g. Th1, Th17) or anti- (T_reg_) inflammatory subsets (1–5). These pro- and anti- inflammatory forces must be precisely balanced; dysregulation of this balance can predispose to autoimmunity, infection or cancer (3, 6).

We have previously demonstrated how small molecules can highlight novel pathways of immunoregulation in primary immune cells. For example, we showed that small molecule inhibition of the dual-specificity tyrosine phosphorylation-regulated kinase 1A (DYRK1A) promotes differentiation of murine and human CD4^+^ T cells into T_reg_s (7). We also showed that small molecule inhibition of salt-induced kinases (SIKs) enhanced production of IL-10 by murine and human myeloid cells (8). However, a comprehensive understanding of how both innate and adaptive immune cell function is modulated remains lacking for most small molecules.

Here, we investigate the effect of the natural product-derived small molecule dihydro- cortistatin A (DCA) on murine and human CD4^+^ T cells. Recent studies pointing to DCA as the CDK8 inhibitor with highest specificity and lowest toxicity highlight DCA as a CDK8 inhibitor of critical interest (*9*). CDK8 is an essential component of the CDK8 submodule of the Mediator coactivator complex, which regulates RNA polymerase II activity (*10, 11*). The CDK8 submodule facultatively binds the Mediator complex, phosphorylates transcription factors and regulates specific pathways (*11*-*13*). CDK8 phosphorylates several immune-relevant transcription factors, including STAT1^Ser727^, STAT3^Ser727^, STAT5^Ser730^, c-Jun^Ser243^ and Notch (*14*-*19*). CDK8 regulates both innate and adaptive immune responses and CDK8 inhibition typically exerts tolerogenic effects. We previously found that DCA promotes production of IL-10 in myeloid cells by inhibiting cyclin-dependent kinase 8 (CDK8) (*20*-*22*). Additionally, CDK8 deletion in innate immune NK cells improves tumor surveillance while in adaptive immune cells, CDK8/19 inhibitors promote T_reg_ differentiation (*18, 22*-*25*). Recent findings that CDK8 inhibition promotes Th17 differentiation suggest the first pro-inflammatory sequelae (*26*). How CDK8 regulates differentiation to other T cell lineages (Th1 and Th2) remains less clear. Furthermore, much of the mechanistic work in T cells has focused on CDK8 phosphorylation of STAT5 and STAT3. The possibility of additional CDK8-regulated pathways in the context of T cell biology is suggested by our findings that CDK8 regulates myeloid cells by c-Jun^Ser243^ phosphorylation; however, the identity of these pathways remains incompletely elucidated (*22*). Understanding these pathways is essential to identify the patients who might most benefit from CDK8 inhibition therapy.

We demonstrate that DCA exerts a unique pattern of immunomodulation (i.e. chemical immunophenotype) compared to other known immunomodulatory small molecules. Using both small molecule inhibitors and CRISPR/Cas9 knockdown, we find that DCA inhibits CDK8 to promote the differentiation of both T_reg_ and Th2 cells while suppressing the differentiation of pro- inflammatory Th1 and Th17 subsets. We show that DCA-driven T_reg_s are fully suppressive in the absence of concomitant tolerogenic effects on innate immune cells. Mechanistically, CDK8 inhibition by DCA regulates T_reg_/Th17/Th1 differentiation independent of effects on STAT1/STAT3 Ser727 phosphorylation. Notably, DCA’s unusual chemical immunophenotype directly leads us to find that DCA uniquely drives early temporal expression of *FOXP3* at least in part via a CDK8-GATA3-FOXP3 pathway not previously described to regulate T_reg_ differentiation. These findings further our mechanistic understanding of an emerging role for DCA as an immunomodulator that broadly drives tolerogenic programs in both innate and adaptive immune cells. These findings are discussed in the context of implications to future therapeutic use of CDK8 inhibitors.

## Results

### DCA exerts tolerogenic effects on murine and human CD4^+^ T cell differentiation

Given our previous observation that DCA promotes tolerogenic IL-10 production in innate immune cells, we determined whether DCA exerts tolerogenic effects on CD4^+^ T cell differentiation (*22*). We tested the effect of DCA on naïve murine CD4^+^ T cells cultured in suboptimal pro-T_reg_ or -Th2 conditions (T_reg_^low^ and Th2^low^, respectively) as we previously described (7). DCA enhanced differentiation of both T_reg_ and Th2 cells (Fig. 1A). DCA increased T_reg_s specifically in cultures of FACS-sorted naïve CD4^+^ T cells, but not sorted T_reg_s, further demonstrating that the increase in T_reg_s is due to enhanced differentiation of T_reg_s rather than expansion of existing T_reg_s (Fig. S1A). To examine if these tolerogenic effects extended to inhibiting differentiation of pro-inflammatory T cell lineages, we added DCA to murine T cells cultured in near-optimal pro-Th1 and -Th17 conditions (Th1^hi^ and Th17^hi^, respectively). DCA significantly inhibited differentiation of Th1 and Th17 cells (Fig. 1B). Notably, DCA promoted differentiation of T_reg_ and Th2 cells even in near-optimal Th17^hi^ and Th1^hi^ conditions respectively (Fig. 1B, FACS plots). In the context of non-polarizing Th0 conditions, DCA significantly, albeit modestly, enhanced murine T_reg_ and Th2 differentiation (Fig. 1C). Th1 differentiation was slightly reduced below the level of statistical significance and Th17 cells were too infrequent to accurately assess (Fig. 1C). These results suggest that DCA can enhance T_reg_/Th2 differentiation in the absence of exogenous cytokines. Therefore, DCA exerts powerful and broad tolerogenic effects on murine T cell differentiation.

**Fig. 1.**
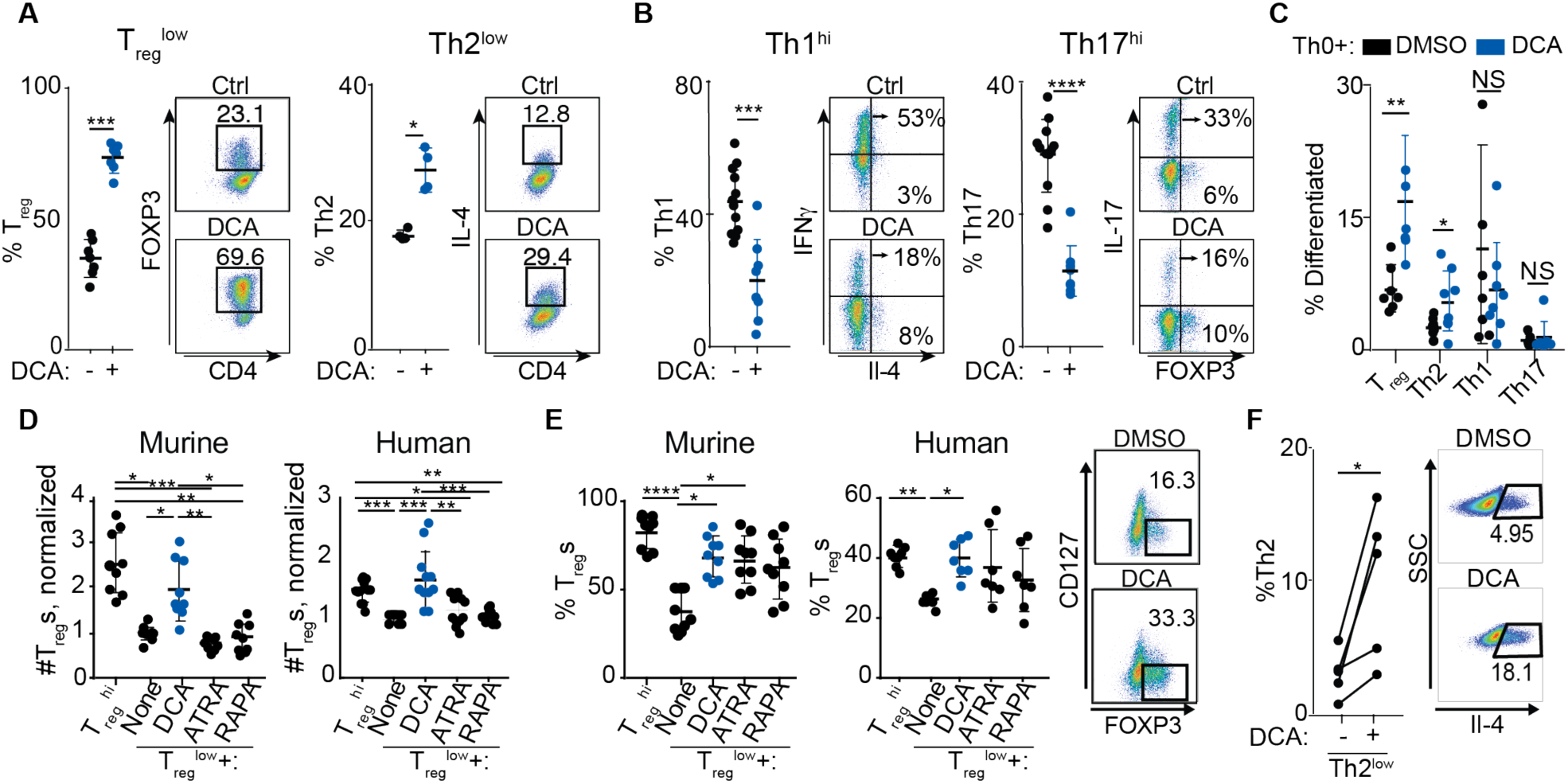
DCA broadly regulates differentiation of murine and human T cells. (A-C) Effect of DCA on murine naïve CD4^+^ T cells cultured in (A) suboptimal pro-T_reg_ or -Th2 conditions (T_reg_^low^ and Th2^low^ respectively), (B) near-optimal pro-Th1 or -Th17 conditions (Th1^hi^ and Th17^hi^ respectively) or (C) neutral Th0 conditions (n = 4-12, x4 experiments). (D-E) Effect of DCA, all- trans retinoic acid (ATRA) and rapamycin (RAPA) on number (D) and percent (E) of T_reg_s generated from murine (n = 9, x4 experiments) and human (n = 7-8, x3 experiments) naïve CD4^+^ T cells cultured in T_reg_^low^ conditions. (F) Effect of DCA on human naïve CD4^+^ T cells cultured in Th2^low^ conditions (n = 5, x2 experiments). Mann-Whitney (A-B), Kruskal-Wallis (C-E) and paired t-test (F) results, *P<0.05, ** P<0.01, *** P<0.001 ****P<0.0001.

We next investigated whether DCA similarly affects human T_reg_ and Th2 differentiation by culturing human CD4^+^ T cells in (human-specific) suboptimal T_reg_^low^ and Th2^low^ conditions respectively. DCA treatment enhanced both T_reg_ and Th2 differentiation in human CD4^+^ T cells, pointing to concordant regulation in human and murine cells (Fig. 1D-F). We benchmarked the pro-T_reg_ effect of DCA against the well-described T_reg_ enhancers all-trans retinoic acid (ATRA) and rapamycin (RAPA) (*27*-*33*). In murine and human CD4^+^ T cells cultured in suboptimal T_reg_^low^ conditions, DCA treatment enhanced the total number of T_reg_s significantly higher than either ATRA or rapamycin (Fig. 1D). In addition, DCA enhanced the percentage of T_reg_s to a level similar to ATRA and rapamycin (Fig. 1E). These results highlight that DCA potently enhances T_reg_ differentiation in both murine and human T cells and reflect in part the lower cytotoxicity of DCA compared to ATRA and rapamycin (Fig. S1B).

### DCA identifies a novel CDK8 inhibition-driven chemical immunophenotype

We compared DCA’s profile of tolerogenic effects against that of other tolerogenic compounds. We investigated the dose-response of murine CD4^+^ T cells to DCA and two other tolerogenic small molecules in the context of suboptimal pro-T_reg_, Th2, Th1 and Th17 conditions (T_reg_^low^, Th2^low^, Th1^low^ and Th17^low^, respectively) (7). These experiments showed that DCA enhanced differentiation of both murine T_reg_ and Th2 cells with identical EC_50_ (dose exerting half- maximal effect), supporting the involvement of a common mechanistic target (Fig. 2A). We wanted to understand if DCA’s ability to enhance T_reg_ and Th2 differentiation and myeloid IL-10 production represent a pattern common to many tolerogenic small molecules. We tested harmine, which we previously identified as a potent enhancer of T_reg_ differentiation, and found that harmine enhanced the differentiation of T_reg_, but not Th2, cells (Fig. 2A) (7). We also tested HG- 9-91-01, which we previously showed enhances myeloid cell production of IL-10 production by inhibiting salt-inducible kinase (SIK) 1-3, and found that HG-9-91-01 enhanced neither T_reg_ nor Th2 differentiation (Fig. 2A) (8). Therefore, DCA, HG-9-91-01 and harmine exert distinct immune phenotypic profiles, which we term chemical immunophenotypes, reflecting engagement of distinct pathways regulating tolerogenicity in innate and adaptive immune cells.

**Fig. 2.**
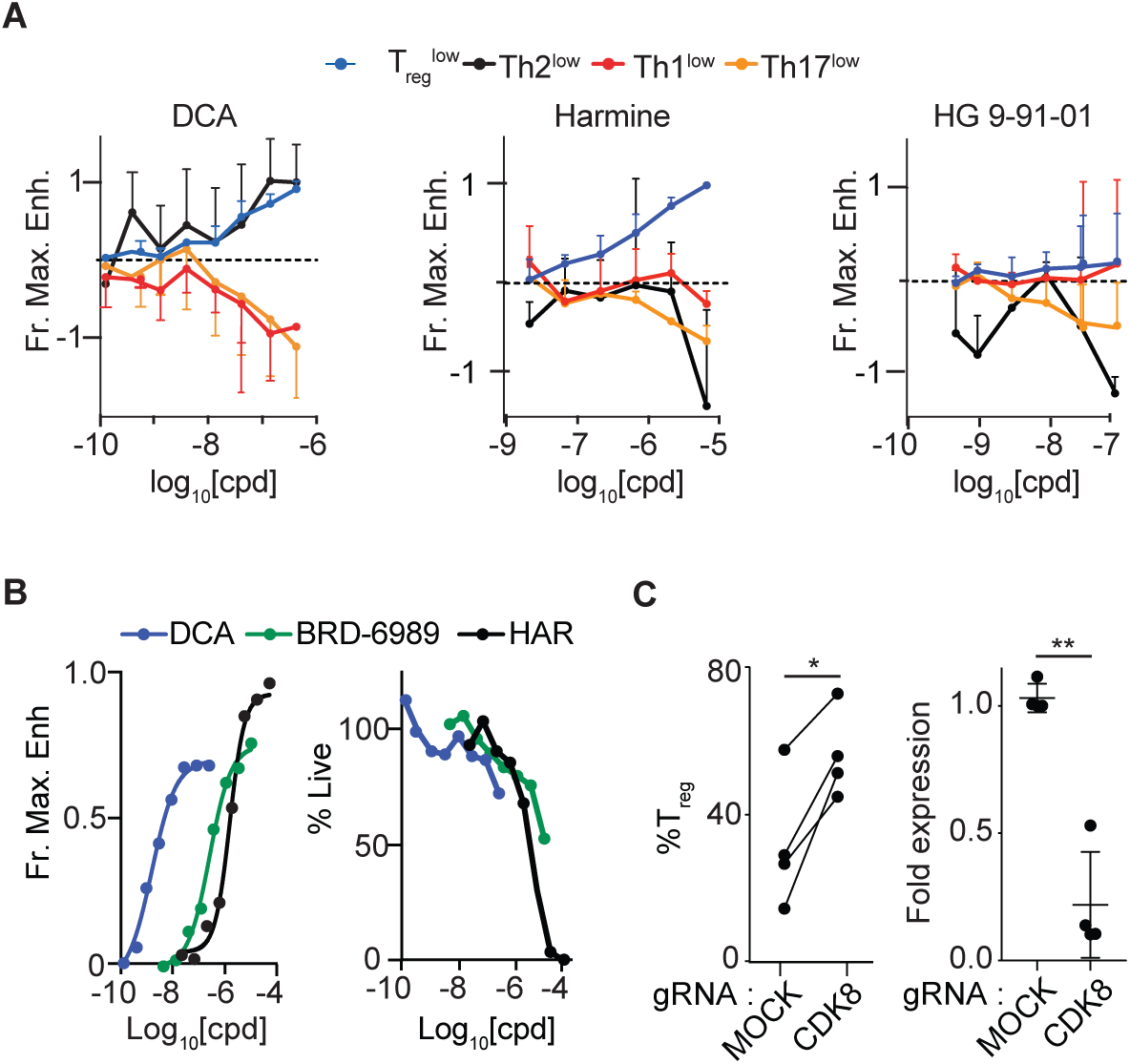
DCA describes a unique chemical immunophenotype. (A) Dose-response curves showing effect of DCA, harmine and HG-9-91-01 on murine CD4^+^ T cells cultured in suboptimal T_reg_^low^, Th2^low^, Th1^low^ and Th17^low^ conditions (n = 3-5, x3-5 experiments). Fractional maximal enhancement was determined by increase in percentage lineage-committed cells, relative to maximal cytokine-driven enhancement as previously reported (7). (B) Naive murine CD4^+^ T cell cultures showing dose-response of the CDK8 inhibitors DCA and BRD-6989 on T_reg_ differentiation (left) and culture cellularity (right) (n = 2, x2 experiments). Harmine (HAR) is included for comparison. (C) Effect of CRISPR/Cas9-mediated deletion of CDK8, compared to mock (no guide) control, on propensity of human CD4^+^ T cells to differentiate into T_reg_s (left) and CDK8 expression (right). (n = 4, x2 experiments). Paired t-test (C), * P<0.05, ** P<0.01.

We and others have previously shown immunomodulatory effects of DCA and other CDK8 inhibitors (*18, 22*-*25*). We used two different approaches to validate CDK8 as the T_reg_- relevant mechanistic target of DCA. Firstly, we tested DCA alongside a structurally distinct small molecule CDK8 inhibitor, BRD-6989 (*22*). In T_reg_^low^ conditions, both CDK8 inhibitors showed concentration-dependent enhancement of murine T_reg_ differentiation with EC_50_ for each compound similar to that observed for enhancing IL-10 production in BMDCs (Fig. 2B) (*22*). The EC_50_ of DCA was much lower than of BRD-6989, driving its subsequent preferential use (Fig. 2B). Notably, DCA and BRD-6989 both exhibited low cytotoxicity, even less than that observed with harmine, which we previously identified as one of the least cytotoxic T_reg_ enhancers (Fig. 2B) (7). Secondly, we used CRISPR/Cas9 to knock out *CDK8* in primary human CD4^+^ T cells. Efficient editing of CDK8 led to enhanced T_reg_ differentiation comparable to levels observed using DCA treatment (Fig. 1E and 2C). These results indicate that DCA enhances murine and human T_reg_ differentiation at least in part by inhibiting CDK8.

### DCA-driven T_reg_s are fully tolerogenic in the absence of DCA-innate immune tolerogenic effects

We next interrogated the suppressive capacity of DCA-driven T_reg_ cells both in vitro and in vivo. Using a standard in vitro suppression assay, we observed no significant differences in the ability of T_reg_^hi^- or T_reg_^low+DCA^-driven murine T_reg_ cells to suppress proliferation of co-cultured responder CD4^+^ T cells (Fig. 3A, red and blue lines respectively). We tested the capacity of DCA- driven T_reg_s to inhibit inflammation in vivo in two murine T_reg_-transfer models in order to exclude confounding effects of systemically-delivered DCA on endogenous innate immune cells. In an established model of type 1 diabetes, transfer of NOD*-BDC2.5^+^* CD4^+^ T cells, specific for an epitope derived from the islet antigen chromogranin A, into NOD*-scid* recipients results in islet β- cell destruction and onset of diabetes about 10 days later (Fig. 3B, black line) (*34, 35*). Co-injection of antigen-specific T_reg_ cells, generated from naïve NOD*-BDC2.5.Foxp3^IRES-GFP^* CD4^+^ T cells using either T_reg_^low+DCA^ or T_reg_^hi^ conditions, significantly delayed onset of diabetes to a similar degree (Fig. 3B, blue and red lines respectively) (7). We observed similar results in a murine model of intestinal inflammation where transfer of CD45RB^hi^CD4^+^ T cells into B10.RAG2^-/-^ recipients results in expansion of donor T cells and inflammation most prominent in the colon about 4 weeks later (Fig. 3C, black) (*36, 37*). Co-transfer of T_reg_ cells, generated from naïve wild-type C57Bl/6 CD4^+^ T cells using either T_reg_^low+DCA^ or T_reg_^hi^ conditions, resulted in significant and similar attenuation of intestinal inflammation (Fig. 3C, blue and red respectively) (*38*). Together, these results demonstrate that DCA-driven T_reg_ cells are fully functional and equivalent to T_reg_^hi^-generated T_reg_ cells (an adoptive cellular therapy-relevant gold standard comparison) both in vitro and in vivo, using model systems employing different genetic backgrounds and T cell specificities. Importantly, these experiments demonstrate that DCA exerts a strong T_reg_-intrinsic tolerogenic effect in the absence of concomitant effects on the innate immune compartment.

**Fig. 3.**
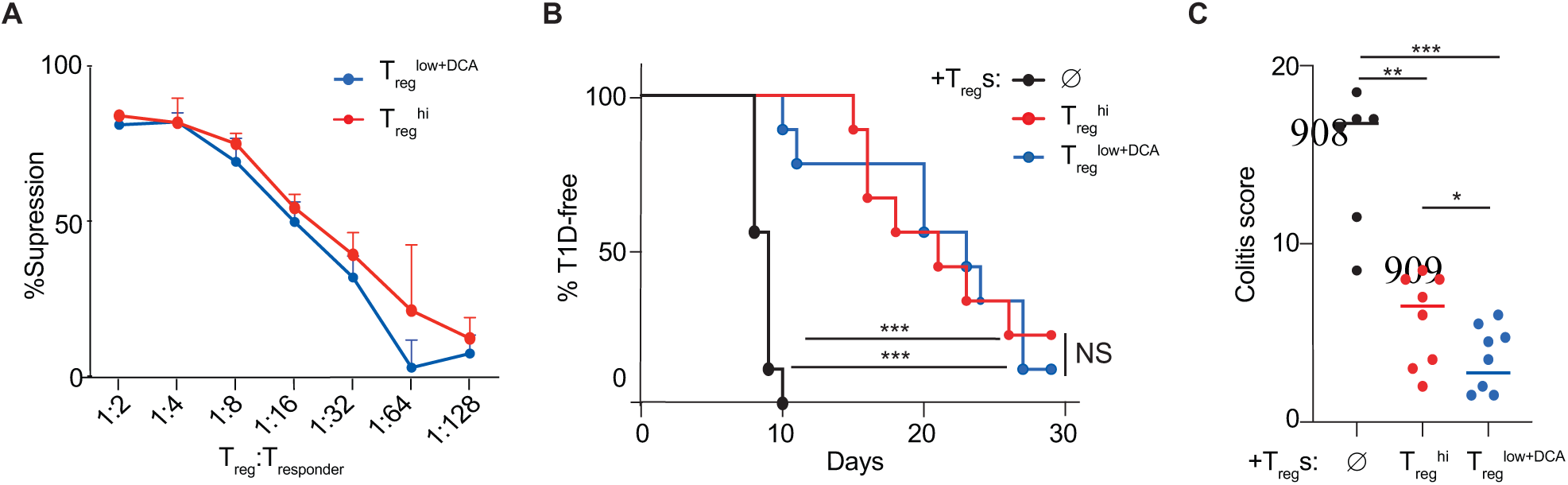
DCA enhances differentiation of functional T_reg_s. Suppressive function of DCA-driven T_reg_s (T_reg_^low+DCA^, blue), compared to T_reg_^hi^-driven T_reg_s (red). (A) Standard in vitro suppression assay, (B) NOD.BDC2.5 model of type 1 diabetes and (C) B10.*Rag2^-/-^* model of colitis. No-T_reg_ controls shown in black. All data representative of at least 2 independent experiments (n≥4 mice per cohort). Mantel-Cox (B) and Mann-Whitney (C) results, * P<0.05, ** P<0.011, *** P<0.001.

### DCA exerts tolerogenic effects on T cell differentiation independent of regulating STAT1^Ser727^, STAT3^Ser727^ and c-Jun^Ser243^ phosphorylation

We investigated key candidates that might account for DCA’s tolerogenic effects in CD4^+^ T cells. CDK8 phosphorylates STAT1^Ser727^ and STAT3^Ser727^ in several cell types (*39*-*42*). Although the role of Ser727 phosphorylation in Th1/Th17/T_reg_ differentiation is unclear, potential contribution is suggested by the central role of STAT1^Tyr701^ and STAT3^Tyr705^ tyrosine phosphorylation in Th1 and Th17 differentiation respectively (*43*-*45*). Recent studies argue that inhibition of CDK8 promotes Th17 differentiation by attenuating STAT3^Ser727^ phosphorylation, emphasizing the importance of investigating this pathway (*26*). DCA reduced IL-6-induced STAT3^Ser727^ phosphorylation in murine CD4^+^ T cells, but did not reduce either STAT3^Tyr705^ phosphorylation or expression of the hallmark Th17 transcription factor RORγt in cells cultured in Th17^hi^ conditions; total STAT3 was slightly decreased (Fig. 4A-C). Primary CD4^+^ T cells from *Stat3^Ser727Ala^* mice, in which Ser727Ala mutation abrogates STAT3^Ser727^ phosphorylation, showed reduced Th17 differentiation, highlighting a previously unappreciated role of STAT3^Ser727^ phosphorylation in this process (Fig. 4D) (*40*). However, DCA suppressed Th17 and enhanced T_reg_ differentiation in both *Stat3^Ser727Ala^* and wild-type CD4^+^ T cells, demonstrating that DCA regulates T_reg_ and Th17 differentiation via mechanisms independent of STAT3^Ser727^ phosphorylation (Fig. 4D-E).

**Fig. 4.**
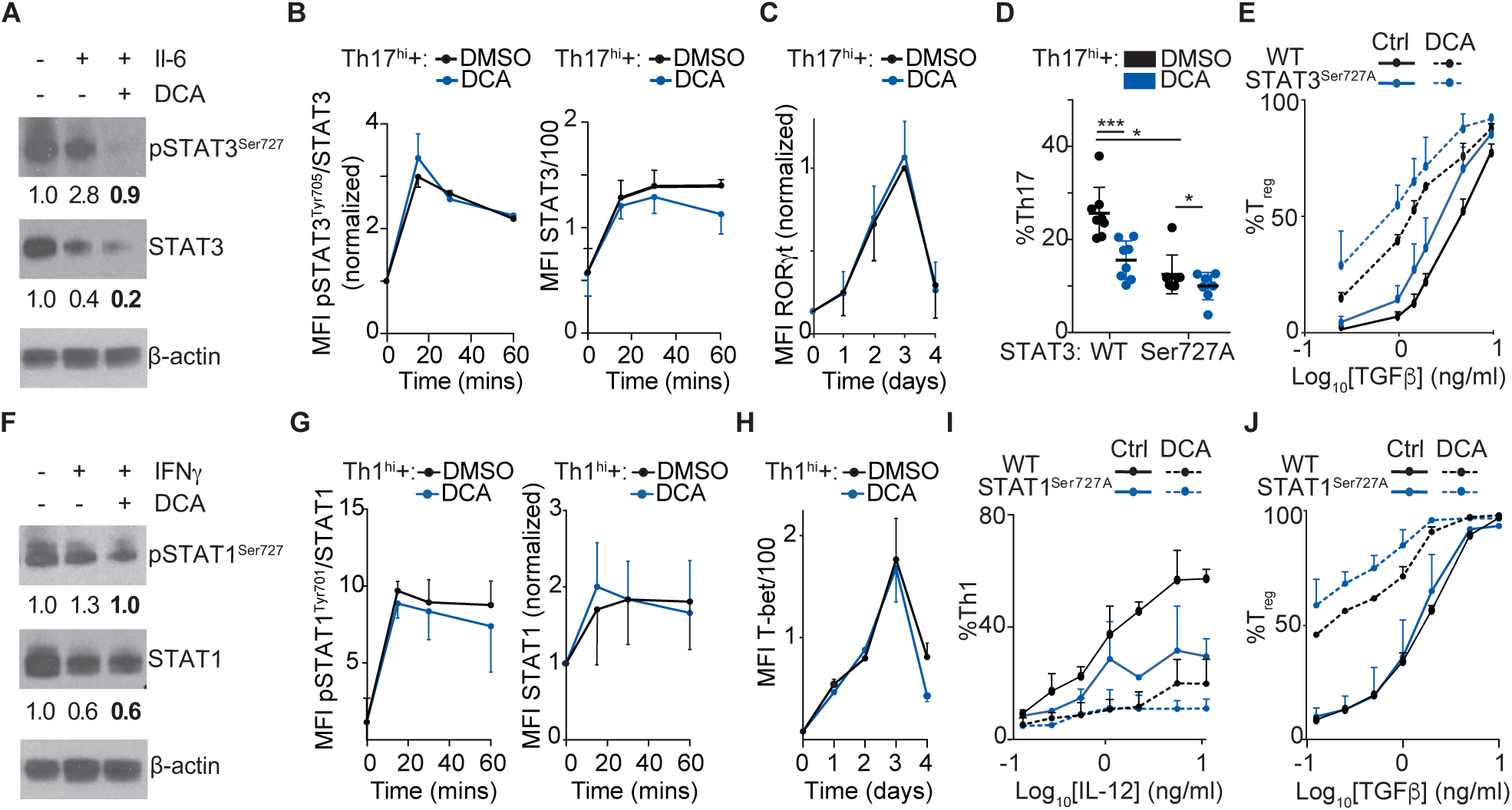
DCA regulates T cell differentiation independently of STAT1/STAT3 Ser727 phosphorylation. (A) Effect of DCA on IL-6-induced STAT3^Ser727^ phosphorylation in resting murine CD4^+^ T cells (representative of 2 independent experiments). (B-C) Effect of DCA on phospho-STAT3^Tyr705^ and total STAT3 (B, n = 2, x2 experiments) and RORγt (C, n = 3, x3 experiments) in murine CD4^+^ T cells cultured in Th17^hI^ conditions. (D-E) Effect of DCA on Th17 (D) and T_reg_ (E) differentiation in *STAT3^Ser727Ala^* naïve murine CD4^+^ T cells. (n = 8, x3 experiments). (F) Effect of DCA on IFNγ-induced STAT1^Ser727^ phosphorylation in resting murine CD4^+^ T cells (representative of 2 independent experiments). (G-H) Effect of DCA on phospho- STAT1^Tyr705^ and total STAT1 (G, n = 2, x2 experiments) and T-bet (H, n = 3, x3 experiments) in murine CD4^+^ T cells cultured in Th1^hI^ conditions. (I-J) Effect of DCA on Th1 (I) and T_reg_ (J) differentiation in *STAT1^Ser727Ala^* naïve murine CD4^+^ T cells (n = 4, x2 experiments). Mann- Whitney * P<0.05, ** P<0.01, *** P<0.001.

We also showed that DCA reduced IFNγ-induced phospho-STAT1^Ser727^ but not of phospho-STAT1^Tyr701^ or total STAT1 in cells cultured in Th1^hi^ conditions (Fig. 4F-G). Expression of the hallmark Th1 transcription factor Tbet in Th1^hi^ cultures was also unaltered by DCA except at day 4, arguing against altered regulation by STAT1^Ser727^ (Fig. 4H). Primary CD4^+^ T cells from *Stat1^Ser727Ala^* mice, in which Ser727Ala mutation abrogates STAT1^Ser727^ phosphorylation, showed reduced Th1 differentiation, highlighting a previously unappreciated role of STAT1^Ser727^ phosphorylation in this process (Fig. 4I) (*41*). However, DCA suppressed Th1 and enhanced T_reg_ differentiation in both *Stat1^Ser727Ala^* and wild-type CD4^+^ T cells, demonstrating that DCA regulates T_reg_ and Th1 differentiation via mechanisms independent of STAT1^Ser727^ phosphorylation (Fig. 4I- J).

Consistent with previously published findings, we also found that treatment with DCA inhibited IL-2-induced phosphorylation of STAT5b^Ser730^ (Fig. S2A) (*18*). We were unable to find *Stat5b^Ser730Ala^* mice to perform similar studies as with STAT1 and STAT3 above.

We next investigated the overlap between characterized CDK8-regulated pathways in innate and adaptive immune cells. Similar to our findings in myeloid cells, T cells cultured in either T_reg_^low^ or T_reg_^hi^ conditions showed increased phosphorylation of both c-Jun^Ser243^ and c- Jun^Ser63^; DCA specifically inhibited phosphorylation on the inhibitory site c-Jun^Ser243^ (Fig. S2B) (*22*). Unlike in myeloid cells, DCA’s tolerogenic pro-T_reg_ effect in T cells was neither attenuated by the AP-1 inhibitor T-5224 nor enhanced by overexpression of multiple c-Jun family members (c-Jun, JunB or JunD) (Fig. S2C-D). Together, these results indicate that DCA regulation of AP-1 transcription factors drives tolerogenicity in myeloid, but not CD4^+^ T, cells.

### DCA enhances Foxp3 expression by engaging GATA3

Temporal flow cytometric analysis of FOXP3 expression throughout the period of culture revealed indistinguishable kinetics between murine CD4^+^ T cells cultured in T_reg_^low^ and T_reg_ conditions until day 2, with FOXP3^+^ cells increasing in T_reg_^hi^ conditions and decreasing in T_reg_^low^ conditions thereafter (Fig. 5A) (7). Notably, DCA treatment significantly increased FOXP3^+^ cells, as well as *Foxp3* expression, at early time points (days 1 and 2) compared to either T_reg_^low^ or T_reg_ conditions (Fig. 5A-B). DCA treatment also drove concordant regulation of other FOXP3- regulated genes at day 2, including upregulation of *Eos, Helios* and *Cd25* as well as downregulation of *Il2* expression (Fig. 5B) (*46*-*49*). These data suggest that DCA promotes murine T_reg_ differentiation at least in part by enhancing early expression of FOXP3. This induction of key T_reg_ transcription factors did not involve canonical T_reg_ pathways; specifically, DCA neither enhanced SMAD2/SMAD3 nor inhibited (mTOR pathway members) S6/S6K phosphorylation, pointing to the involvement of novel pathway(s) (Fig. S3A-B).

**Fig. 5.**
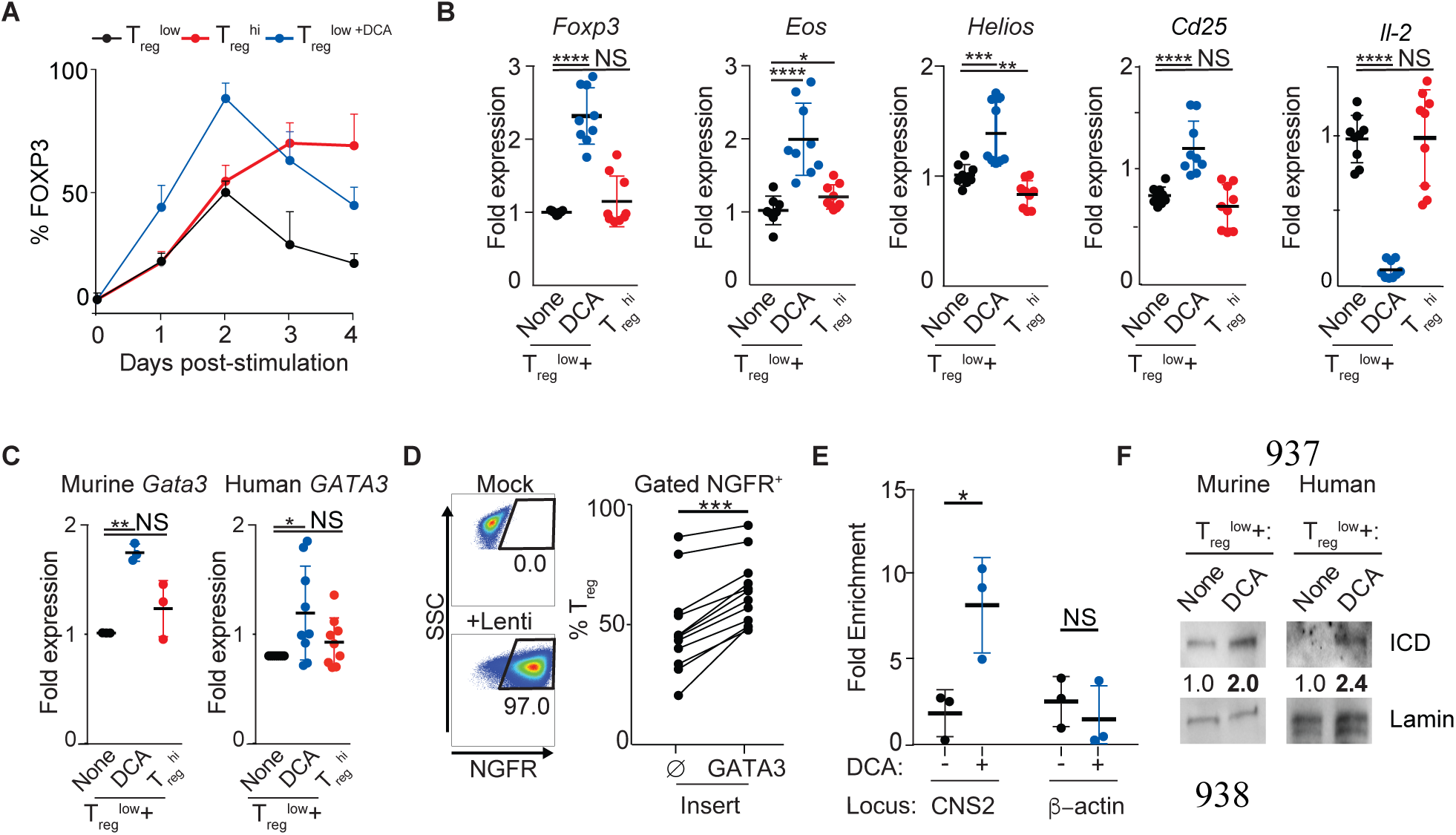
DCA drives novel early FOXP3 expression via a novel CDK8-Notch-GATA3 pathway. (A) Timecourse of FOXP3 expression in murine CD4^+^ T cells cultured in T_reg_^low^, T_reg_^low+DCA^ and T_reg_ conditions (n = 10, x5 experiments). (B) Effect of DCA on expression of FOXP3-regulated genes in murine CD4^+^ T cells cultured for 2 days in T_reg_^low^ conditions (n = 9, x3 experiments). (C) Effect of DCA on *Gata3* expression in murine (n = 3, x3 experiments) and human (n = 9, x3 experiments) CD4^+^ T cells, cultured for 2 days in T_reg_^low^ conditions. (D) Effect of overexpressing GATA3, using transduction of either NGFR-T2A-GATA3 or NGFR control lentivirus, on T_reg_ differentiation in human CD4^+^ T cells cultured in T_reg_^low^ conditions (n = 12, x3 experiments). (E) ChIP-qPCR quantitation of how DCA treatment impacts GATA3 binding to FOXP3 CNS2 in human CD4^+^ T cells, cultured for 2 days in T_reg_^low^ conditions (n = 3, x3 experiments). β-actin is included as a control locus. (F) Effect of DCA on intranuclear levels of Notch intracellular domain (ICD), normalized to nuclear Lamin B1 levels, in murine and human CD4^+^ T cells stimulated for 2 hours in indicated conditions (representative of ≥2 independent experiments). Mann-Whitney (B), Wilcoxan matched pair analysis (C-D) and paired t-test (E), * P<0.05, ** P<0.01, *** P<0.001 ****P<0.0001.

DCA’s unusual chemical immunophenotype led us to consider a mechanistic link between DCA-mediated enhancement of both T_reg_ and Th2 differentiation. GATA3, the hallmark Th2 transcription factor, is highly expressed beginning at the earliest timepoints of Th2 differentiation (*50*). Although not well studied in the context of T_reg_ differentiation, previous findings that GATA3 binds the CNS2 enhancer element of the *FOXP3* locus and regulates mature T_reg_ physiology support the possibility that GATA3 could regulate FOXP3 expression and thus T_reg_ differentiation (*51*-*53*). Consistent with this notion, DCA treatment of both murine and human CD4^+^ T cells cultured in T_reg_^low^ conditions also enhanced early *GATA3* expression at day 2 (Fig. 5C). To validate the role of DCA-mediated upregulation of GATA3 in T_reg_ differentiation, we generated lentiviral vectors to overexpress *GATA3*, including a truncated NGFR marker separated by a self-splicing T2A peptide to allow specific comparison of transduced cells. We performed these experiments using human T cells because these were more amenable to viral transduction. Overexpression of *GATA3* consistently enhanced T_reg_ differentiation in human naïve CD4^+^ T cells cultured in T_reg_^low^ conditions, compared to cells transduced with control (NGFR only) virus (Fig. 5D). To further support the functional significance of DCA-enhanced GATA3, we performed chromatin immunoprecipitation experiments and found that DCA treatment of human CD4^+^ T cells cultured in T_reg_^low^ conditions resulted in significantly increased binding of GATA3 specifically to *FOXP3* CNS2 early in T_reg_ differentiation (Fig. 5E). These results argue that GATA3 is an early regulator of *FOXP3* expression and T_reg_ differentiation that can be regulated by DCA.

We sought to better understand how DCA might regulate *GATA3* expression. Previous studies showed that Notch can directly drive *Gata3* expression and that enhanced Notch signaling promotes T_reg_ differentiation (*54, 55*). Furthermore, CDK8 inhibits Notch signaling by phosphorylating the Notch signaling domain ICD1, leading to its degradation, leading us to hypothesize that DCA may drive *Gata3* expression by enhancing Notch signaling in T cells (*19*). Consistent with this notion, we found that treatment with DCA led to increased intranuclear levels of ICD1 in both murine and human CD4^+^ T cells (Fig. 5F). Supporting functional relevance of DCA-driven increased intracellular ICD1, we performed RNAseq analyses comparing FOXP3^+^ cells cultured for 2 days in T_reg_^low^ versus T_reg_^low+DCA^ conditions to define a 577-gene signature associated with DCA treatment. Using a previously described method, we showed that this signature is maintained in sorted mature FOXP3^+^ iT_reg_s generated after 4 days of culture in T_reg_^low+DCA^ versus T_reg_^hi^ conditions and partially maintained in FOXP3^+^ versus FOXP3^-^ cells cultured for 2 days in T_reg_^low^ conditions, supporting FOXP3- and T_reg_-relevance of this signature (Fig S4) (*7, 56*). Transcription factor target analysis of this DCA signature using the ChIP-seq result-based Gene Transcription Regulation Database (GTRD) in GSEA revealed MAML as the most enriched transcription factor (Table S1) (*57*-*59*). MAML is recruited by DNA-bound ICD1, in complex with RBP-J, whereupon it recruits transcriptional co-activators (*60*). Therefore, enrichment of MAML binding sites is consistent with enrichment of ICD1 binding to DCA- regulated genes. Together, these results demonstrate for the first time that inhibition of CDK8 by DCA drives *FOXP3* expression and T_reg_ differentiation at least in part by driving increased Notch and GATA3 signaling.

## Discussion

Here we have demonstrated that DCA promotes T_reg_ differentiation at least in part by engaging a previously unappreciated CDK8-GATA3-FOXP3 pathway. Our use of both novel small molecules (DCA and BRD-6989) and CRISPR/Cas9-mediated deletion point to CDK8 inhibition as the mechanism by which DCA exerts these effects. While previous studies have shown that GATA3 impacts mature T_reg_ physiology, this is the first report to our knowledge that GATA3 can drive T_reg_ differentiation (*51, 61*). Our findings extend prior studies showing that GATA3 protein levels are upregulated during T_reg_ differentiation (*62*). Whether GATA3 initiates, stabilizes or amplifies *FOXP3* expression remains to be dissected in future studies. Previous reports suggesting that GATA3 inhibits FOXP3 expression used significantly different experimental approaches, including secondary CD3 stimulation or removal of primary TCR stimulation, and did not exclude contribution of IL-4 (*63, 64*). Given that GATA3 is a hallmark Th2 transcription factor that inhibits Th1 and likely Th17 differentiation, this CDK8-GATA3- FOXP3 pathway provides a parsimonious unifying mechanism to explain, at least in part, how DCA broadly regulates T_reg_, Th2, Th1 and Th17 differentiation (*65*-*67*). Given that Th2 cells can produce IL-10, this previously unappreciated link between T_reg_ and Th2 differentiation may point to conserved (e.g. CDK8-related) anti-inflammatory signaling pathways that can engage distinct downstream effector pathways (*68*). We hypothesize that differences in the local cytokine milieu impact whether DCA enhances T_reg_ or Th2 differentiation, for example by modulating epigenetic accessibility of *FOXP3* and *IL4* loci. Additionally, where CDK8 effector pathways regulating tolerogenicity in innate (via phospho-c-Jun^Ser243^) vs adaptive immune cells diverge remains to be clearly defined (*22*).

Our discovery of DCA’s unusual temporal profile of enhancing early expression FOXP3 and many FOXP3-regulated genes contrasts with the temporal profile of T_reg_^hi^ conditions and other T_reg_-enhancing compounds like harmine and reinforces a model of T_reg_ differentiation that involves independently regulated early and late pathways (7). Whereas early pathways might involve TGFβ licensing cells to adopt T_reg_ fate and express FOXP3, late pathways might maintain and promote T_reg_ lineage commitment. Our data support a model where DCA largely enhances early pathways, including Notch-GATA3, to regulate FOXP3 expression. This suggests DCA may have particular therapeutic relevance to patients who have defects in early pathways of T_reg_ differentiation and also raises the possibility of synergy with therapies that enhance late pathways of T_reg_ differentiation.

Our findings exemplify how chemical immunotypes point to an important classification scheme that can inform both mechanistic and therapeutic hypotheses. DCA’s unique chemical immunophenotype (pro-T_reg_, pro-Th2 and pro-myeloid-IL-10) is distinct from that of many other tolerogenic compounds, including SIK- and DYRK1A-inhibitors, which exert tolerogenic effects specifically in either innate or adaptive immune cells, but not both (*7, 22*). Our novel finding that CDK8 inhibition promotes both T_reg_ and Th2 differentiation directly informed our interrogation of GATA3 as a putative regulator of T_reg_ differentiation. Additionally, our studies suggest value in monitoring tolerogenic effects, including impaired host-versus-tumor effects, in anticipated clinical use of CDK8 inhibitors as cancer therapeutics (*69, 70*).

The translational relevance of these data is reinforced by our finding that DCA promotes T_reg_ differentiation in primary human CD4^+^ T cells. We note that T_reg_s generated using DCA are fully functional in vitro and in vivo. Importantly, our use of T_reg_-transfer models specifically interrogates the functionality of DCA-driven T_reg_s without confounding anti-inflammatory effects on innate immune cells, that could confound the interpretation of models using systemic drug administration (*18, 25*). DCA and other CDK8 inhibitors may find utility as tolerogenic immunomodulators. Studies suggesting poor long-term tolerability of CDK8 inhibitors, together with our data showing DCA impacts early pathways in T_reg_ differentiation, support this consideration (*71*). We recognize the utility of DCA in generating T_reg_s ex vivo, which would circumvent concerns regarding toxicity in vivo (*71*).

Our findings highlight the value of definitive interrogation of regulatory pathways. Prior knowledge drove the notion of CDK8-STAT interactions as key candidates to explain how CDK8 inhibition regulates T cell differentiation (*14*). Our experiments using *Stat1^Ser727Ala^* and *Stat3^Ser727Ala^* mice clearly demonstrate that DCA regulates Th1, Th17 and T_reg_ differentiation independent of its effects on STAT1^Ser727^/STAT3^Ser727^ phosphorylation. Prior studies suggest that STAT1^Ser727^/STAT3^Ser727^ phosphorylation is required for full transcriptional activity (*39*-*42*). Consistent with this, we demonstrate a previously unappreciated role of STAT1^Ser727^ and STAT3^Ser727^ phosphorylation in positively regulating Th1 and Th17 differentiation respectively. These findings differ from recent studies suggesting that CDK8 inhibition promotes human Th17 differentiation; possible explanations include differences in species, CDK8 inhibitor and experimental approach (knockin versus transduced allele) (*26*). This emphasizes the value of Ser-Ala STAT mutant mice in dissecting (CDK8-related) mechanistic hypotheses, including developing *Stat5b^Ser730Ala^* mice to definitively define the role of CDK8-regulated STAT5b^Ser730^ phosphorylation in T_reg_ differentiation (*18*). These findings have important implications for disease pathobiology and precision therapy, for example suggesting synergy of therapeutically targeting STAT1^Tyr701^/STAT3^Tyr705^, STAT1^Ser727^/STAT3^Ser727^ and CDK8.

In summary, our studies highlight CDK8 as a regulator of innate and adaptive immune tolerogenicity that is therapeutically targeted by the high-specificity low-toxicity inhibitor DCA (*9*). We show for the first time that CDK8 regulates Th2 differentiation, and human T_reg_ differentiation. The unique chemical immunophenotype of DCA (pro-T_reg_/Th2) directly informs the discovery of a novel CDK8-Notch-GATA3-FOXP3 axis that regulates early pathways of T_reg_ differentiation and has further mechanistic and therapeutic implications. Our demonstration that DCA effectively enhances T_reg_ differentiation compared to canonical T_reg_ enhancers suggests utility in approaches to generate T_reg_s ex vivo for adoptive cellular therapy. In addition, the broadly tolerogenic effects of DCA suggest that it may broadly be useful in the setting of pathologic inflammation or autoimmunity.

## Materials and Methods

### Mice

Balb/c RRID:IMSR_JAX:000651, C57Bl/6 000664RRID:IMSR_JAX:000664, *Foxp3^IRES-GFP^* RRID:IMSR_JAX:006772*, CD45.1^+/+^*002014RRID:IMSR_JAX:002014, NOD-*scid* and NOD-*BDC2.5* mice were purchased from Jackson Labs. NOD-*BDC2.5.Foxp3^IRES-GFP^* mice were from the JDRF Transgenic Core (Harvard Medical School, Boston, MA). C57Bl/10-*Rag2^−/−^* mice were a kind gift from Brian Kelsall (*37*). *Stat1^Ser727Ala^* and *Stat3^Ser727Ala^* mice were previously described (*40, 41*). Mice were housed in the Benaroya Research Institute Vivarium in a SPF animal room with unfettered access to food and water. All murine experiments were performed on male and female mice between 7-12 weeks of age, with the approval of the IACUC of Benaroya Research Institute (Seattle, WA).

### Human samples

Frozen PBMCs and fresh peripheral blood samples were obtained from the Benaroya Research Institute Immune Mediated Disease Registry and Repository. Human studies were approved by the Benaroya Research Institute’s Institutional Review Board and all subjects signed written informed consent prior to inclusion in the study.

### Cell lines

293T cells used in lentiviral production were a generous gift from David Rawlings. 293T cells are female. They were cultured DMEM medium (Hyclone) supplemented with fetal bovine serum and glutamax (Thermo fisher) at 37°C and 5% CO2. Cells were split every 3 days at a density of 7.5×10^4 cells per ml.

### Small Molecules and Reagents

Δ16-cortistatin A (DCA) was a generous gift from P. Baran (The Scripps Research Institute) and synthesized as previously reported (*20, 72*). Small-molecule reagents were confirmed to have ≥95% purity by HPLC–MS. Antibodies, chemical reagents and cytokines were sourced as listed in Table S2. Primers are listed in Table S3

### Cloning and Plasmid Preparation

Coding sequences of GATA3, JUND, c-JUN and JUNB were PCR amplified from pHAGE- GATA3, JunD-HA neo, pMIEG3-c-Jun and pMIEG3-JunB (Addgene #116747, 58515, 40348 and 40349, gifts from Gordon Mills & Kenneth Scott, Kevin Janes, and Alexander Dent respectively). PCR overhang extension was used to add (i) self-splicing T2A sequence and (ii) 40 base pair homology arms (HA) to permit cloning into into EcoRV-digested pLKO.NGFR using Gibson assembly ultra-kit (Codex DNA, San Diego, CA). Primers used are listed in Table S2.

### Murine T cell isolation and culture

Unless otherwise noted, CD4^+^ CD62L^+^ naïve T cells were isolated from 8-12 week-old mice using CD4 negative enrichment kits (Stemcell Technologies, Vancouver, Canada) and CD62L microbeads (Miltenyi Biotec, San Diego, CA) according to the manufacturer’s instructions and confirmed >90% pure by flow cytometry. Cells were cultured on 96 well plates pre-coated with anti-CD3 and anti-CD28 using conditions outlined in Table S4. The addition of DCA to T_reg_^low^ conditions is abbreviated as T_reg_^low+DCA^. T_reg_ and Th1 cultures were fed with equal volume of IL-2 supplemented media (20ng/ml) and retreated with compound at day 2, split 1:2 into IL-2- supplemented media (10 ng/ml) at day 3 and analyzed at day 4. Th17 cultures were treated similarly except no IL-2 was supplemented. Th2 cultures were treated similarly as T_reg_ cultures except they were additionally split 1:2 into IL-2 supplemented media (10 ng/ml) at day 4 and day 5 and analyzed on day 6. To assess STAT1/STAT3/STAT5b Ser phosphorylation, cells were stimulated with 10 ng/ml IFNγ + 2μg/ml anti-IL-4, 10 ng/ml IL-6 + 2 μg/ml each anti-IL4/-12/- IFNγ and anti-CD3/CD28 + 100 ng/ml IL-2 respectively.

### Human T cell isolation and culture

Frozen PBMCs and fresh peripheral blood samples were obtained from the Benaroya Research Institute Immune Mediated Disease Registry and Repository. Human peripheral blood mononuclear cells were isolated from fresh whole blood by Ficoll-Paque (GE Healthcare, Little Chalfont, United Kingdom). CD4^+^CD45RA^+^ naïve T cells were isolated using negative enrichment kits (Stemcell Technologies, Vancouver, Canada) per manufacturer’s instructions and confirmed >90% pure by flow cytometry. Cells were cultured on 96 well plates pre-coated with anti-CD3 and anti-CD28 using conditions outlined in Table S4. T_reg_ cultures were fed with equal volume of IL-2 supplemented media (20ng/ml) and retreated with compound at day 2, split 1:2 into IL-2-supplemented media (10 ng/ml) at day 4 and analyzed at day 5. Th2 cultures were fed and split into media supplemented with IL-2+IL-4 (20 ng/ml each at day 2, 10 ng/ml each thereafter) and compound as indicated to maintain ∼10^6^ cells/ml, restimulated on days 7 and 14 on plates pre-coated with anti-CD3 and anti-CD28 and analyzed at day 21 as previously described (*73*).

### Lentiviral Production

On day 0, 3.8 x 10^6^ 293T cells were plated in 10 ml DMEM + 5% Glutamax (Thermofisher) on a 10 cm plate. On day 1, cells were transfected with 1.5 μg pMD2G, 3 μg psPAX2 (kind gifts from David Rawlings) and 6 μg of pLKO vector, mixed with 42 μg PEI transfection reagent (Polysciences, Inc.) and suspended in 0.5 ml diluent (10 mM HEPES, 150mM NaCl, pH 7.05). Cells were PBS-washed and fed with fresh DMEM + Glutamax on day 2. Viral supernatant was harvested on day 4, centrifuged (2000 rpm x 5 mins) to remove cellular debris, overlaid onto 5 ml of 10% sucrose in NTE (135 mM NaCl, 10mM TrisCl, ph 7.50, 1mM EDTA) in ultra-centrifuge tubes (Beckman) and centrifuged at 25,000 rpm for 90 minutes at 4°C. Supernatant was removed and viral pellet resuspended in ice cold NTE by shaking for 2 hours at 4°C.

### RNP complexing

RNPs were generated by mixing 1.25 ug Cas9 protein (Aldevron, Fargo, ND) and 2.5 pmol each of 3 sgRNAs (Synthego, Menlo Park, CA) with gentle swirling, and incubating at 37°C for 15 minutes. Guides used were CDK8: CUCAUGCUGAUAGGAAG, UGUUUCUGUCUCAUGCUGAU, and UCUGUCUCAUGCUGAUAGGA.

### CRISPR-Cas9 gene editing

CRISPR-Cas9 gene editing was performed as previously described with modifications (Aksoy et al., 2020; Roth et al., 2018). Briefly, human CD4^+^CD45RA^+^ naïve T cells were cultured on 96 well plates pre-coated with anti-CD3 and anti-CD28 in TCM supplemented with 5% Fetal Bovine Serum, 20 ng/ml IL-2 and 2 μg/ml each of anti-IL-12, anti-IFNγ and anti-Il-4. Cells were harvested 2 days later, centrifuged (90 g for 8 minutes), resuspended in buffer T, mixed with 20μM of each RNP complex and electroporated (1600 volts, 10 ms, 3 pulses) using a Neon transfection system (Thermo Fisher, Waltham, MA). Cells were transferred into 90 μl TCM pre- warmed to 37°C. After 24 hours, cells were fed with media supplemented with 100 ng/ml IL-2 and 1 ng/ml TGFβ. Cells were maintained for 5 additional days at a density of 1×10^6^/ml and then analyzed by flow cytometry.

### Flow Cytometry

Cells were stimulated with PMA and ionomycin (50 and 500ng/ml respectively) (Sigma Aldrich, St. Louis, MO) in the presence of Golgistop (BD Biosciences, San Jose, CA) 5 hours prior to analysis as necessary. Cells were typically stained with LIVE/DEAD (Thermo Fisher, Waltham, MA) and anti-CD4 prior to fixation and permeabilization, which was generally performed with either Foxp3 fixation/permeabilization buffers (eBioscience, San Diego, CA). Phosflow cell lyse/fix and PermIII buffers (BD Biosciences, San Jose, CA) were used for phospho-protein assessment. Intracellular staining was performed per manufacturer’s instructions. Counting beads (10 µm, Spherotech, Lake Forest, IL) were added at 5000 per sample. Acquisition was performed on either a FACScalibur or a FACScanto (BD Biosciences, San Jose, CA). Cell sorting was performed using a FACs Aria II (BD Biosciences, San Jose, CA). Data was analyzed using FlowJo software (Treestar, Ashland, OR). Fractional maximal enhancement was determined by increase in percentage lineage-committed cells, relative to maximal cytokine-driven enhancement as previously reported (7). Fractional inhibition was calculated relative to DMSO treated cells (7). STAT1/STAT3 phosphorylation was quantified as previously described (*74*).

### In vitro proliferation and T_reg_ suppression assay

These were performed as previously described (*75*). Briefly, sorted CD45.1^+^CD4^+^CD62L^+^ T_responders_ were labeled with CellTrace Far Red (Thermo Fisher, Waltham, MA) per manufacturer’s protocol and plated at 5×10^4^ cells per well in 96-well U-bottom plates in the presence of anti-CD3 anti-CD28 beads (Dynabead, Grand Island, NY). For T_reg_ suppression assays, T_responders_ were co- cultured with sorted CD45.2^+^Foxp3^IRES-GFP+^ T_reg_ cells generated as indicated. Cells were analyzed by flow cytometry 3 days later.

### T_reg_ suppression – Type 1 diabetes model

These were performed as previously described (7). Briefly, 5×10^4^ sorted CD4^+^CD62L^+^ naïve T cells isolated from NOD-*BDC2.5^+^* mice were injected intravenously into NOD-*scid* mice with or without 1×10^5^ T_reg_ cells generated from NOD-*BDC2.5^+^FOXP3^IRES-GFP^* mice as indicated (*34, 35*). Blood glucose levels were monitored with a handheld Contour glucometer (Bayer, Leverkusen, Germany) at days 3, 6, 8 and every day following. Diabetes was diagnosed when blood sugar exceeded 250 mg/dl for 2 consecutive days.

### T_reg_ suppression – CD45RB^hi^ colitis model

As previously described 5×10^5^ sorted CD4^+^CD62L^+^ naïve T cells isolated from CD45.1^+^ mice were injected intravenously into B10-*Rag2^-/-^* mice (*37, 38*). 5 days later, mice were injected with either PBS or 1.5×10^5^ T_reg_ cells generated from Foxp3^IRES-GFP^ mice as indicated (*38*). Mice were monitored at least weekly for weight loss and morbidity per protocol. Mice were euthanized after 4 weeks and proximal, medial, and distal colon analyzed histologically by blinded observers as previously described (*76*).

### Histology

Tissues were preserved in 10% formalin. Paraffin embedding, sectioning and staining with hematoxylin and eosin was performed by the Histology Core (Benaroya Research Institute, Seattle, WA).

### Western Blotting

Cells were washed in PBS and lysed in either TNN lysis buffer, pH 8 (100 mM TRIS-HCl, 100 mM NaCl, 1% NP-40, 1 mM DTT, 10 mM NaF) or RIPA lysis buffer (150 mM NaCl, 1% Triton X-100, 0.5% sodium deoxycholate, 0.1% SDS, 50 mM TRIS-HCl at pH7.8) supplemented with DTT, protease inhibitors (Roche, Indianapolis, IN) and phosphatase inhibitors (Cell Signaling Technologies, Danvers, MA). Lysates were separated by SDS-PAGE using Tris-Glycine gels loaded with about 1×10^6^ cell equivalents per well and transferred onto PDVF membrane (Millipore, Burlington, MA). Blots were blocked in either 5% Milk (Nestle, Vervey, Switzerland) or bovine serum albumin (Sigma Aldrich, St. Louis, MO) and visualized with Western Lightning Plus-ECL (Perkin Elmer, Waltham, MA) and/or SuperSignal West Femto substrate (Thermo Scientific, Waltham, MA) per manufacturer’s instructions. Nuclear isolation was performed using Nuclei EZ Prep kit per manufacturer’s instructions (Sigma Aldrich, St. Louis, MO). Fractions were subsequently lysed with Triton X-100 lysis buffer (1% Triton X-100, 150 mM NaCl, 50 mM Tris-HCl pH7.8). Band intensity was quantified by ImageJ (*77*).

### RNA Isolation and qRT-PCR

RNA was isolated using RNeasy kits (Qiagen, Valencia, CA) and cDNA generated using iScript cDNA synthesis kits (BioRad, Hercules, CA) per manufacturer’s directions. Real-time PCR was performed using an ABI 7500 FAST REAL-TIME PCR (Applied Biosystems, Foster City, CA) system. Cycling conditions were as follows; 1 cycle of 50°C for 2 minutes, 95°C for 10 minutes, followed by 40 cycles of 95°C for 15 seconds, and 60°C for 1 minute. Primers used are listed in Table S3.

### RNA-seq library preparation and sequencing

RNA-seq libraries were generated from four *Foxp3^GFP^* littermate mice. On day 0, 1000 naïve CD4^+^CD62L^+^ cells were sorted for RNA-seq. The remaining cells were cultured on plates pre-coated with anti-CD3 and anti-CD28 in T_reg_^low^, T_reg_^hi^ and T_reg_^low+DCA^ conditions. On day 2, 250 FOXP3^+^ cells and 500 FOXP3^-^ cells were sorted from cells cultured in T_reg_^low^, T_reg_^hi^ and T_reg_^low+DCA^ conditions. On day 4, 1000 FOXP3^+^ cells were sorted from T_reg_^hi^ and T_reg_^low+DCA^ cultures. Cells were sorted directly into lysis buffer from the SMART-Seq v4 Ultra Low Input RNA Kit for Sequencing (Takara) and frozen until all samples were ready for simultaneous processing. Reverse transcription was performed followed by PCR amplification to generate full length amplified cDNA. Sequencing libraries were constructed using the NexteraXT DNA sample preparation kit (Illumina) to generate Illumina-compatible barcoded libraries. Libraries were pooled and quantified using a Qubit® Fluorometer (Life Technologies). Dual-index, single-read sequencing of pooled libraries was carried out on a HiSeq2500 sequencer (Illumina) with 58-base reads, using HiSeq v4 Cluster and SBS kits (Illumina) with a target depth of 5 million reads per sample.

Base-calling was performed automatically by Illumina real time analysis software. Demultiplexing to generate FASTQ files was performed by bcl2fastq running on the Illumina BaseSpace platform. Subsequent processing was performed using the Galaxy platform. FASTQ reads were trimmed in two steps: 1) hard-trimming to remove 1 3’-end base (FASTQ Trimmer tool, v.1.0.0); 2) quality trimming from both ends until minimum base quality for each read ≥ 30 (FastqMcf, v.1.1.2). Reads were aligned to the GRCm38 mouse reference genome using STAR v.2.4.2a, with gene annotations from GRCm38 Ensembl release number 91 (*78*). Read counts per Ensembl gene ID were quantified using htseq-count v.0.4.1 (*79*). Sequencing, alignment, and quantitation metrics were obtained for FASTQ, BAM/SAM, and count files in Galaxy using FastQC v0.11.3, Picard v1.128, Samtools v1.2, and htseq-count v.0.4.1 (*80*). RNAseq data were then processed using Tidyverse, Biomart, EdgeR and limma to generate relative expression values (*81*-*84*). The raw RNA-seq data has been deposited to the Gene Expression Omnibus (GEO) with accession number GSE141933.

### Pathway Analysis

Pathway analysis was performed using the Gene Set Enrichment Analysis Molecular Signature Database or MSigDB v7.0 which uses the hypergeometric distribution on a background of all genes to calculate a p-value (*58, 85, 86*).

### Microscale Chromatin Immunoprecipitation Assay

Assay was performed as described previously with few modifications (*87*). 100,000 naïve CD4^+^ T-cells were cultured in T_reg_^low^ and T_reg_^low+DCA^ conditions for 2 days and then harvested, washed with ice cold PBS, fixed using 10% v/v of 11% formaldehyde (diluted from 36% stock in 50mM HEPES pH 7.5, 100 mm NaCl, 1 mM EDTA, 0.5 mM EGTA) for 10 minutes, quenched using 5% v/v 2.5 M glycine for 5 minutes, washed twice with 1ml ice-cold PBS and lysed in 50 μl lysis buffer (50mM Tris-HCL pH 8.0, 10 mM EDTA, 1%SDS, 20mg/ml sodium butyrate) supplemented with phenyl methane sulfonyl fluoride and protease inhibitor cocktail (Active Motif, Carlsbad, CA). DNA was sheared by sonication (Biorupter, Diagenode, Denville, NJ) into 200-500 bp fragments. Chromatin was pre-cleared using 30 μl protein G agarose beads (Active Motif) pre-blocked with BSA per manufacturer’s instructions; beads were then removed by centrifugation. Chromatin was diluted with equal volume PBS, 4 μl of anti-GATA3 or isotype control (Cell Signaling Technologies, Danvers, MA) added and sample incubated at 4°C with end over end rotation. Next, 30 μl of pre-blocked Protein G Agarose beads was added and sample incubated for 4 hours at 4°C. Beads were then sequentially washed with 1 ml each low SDS lysis buffer (50mM Tris-HCL pH 8.0, 10 mM EDTA, 0.1%SDS, 20mg/ml sodium butyrate), low salt buffer (10 mM Tris-HCl, pH 8, 1 mM EDTA, 50 mM NaCl), high salt buffer (50 mM Tris-HCl, pH 8, 500 mM NaCl, 0.1% SDS, 0.5% Na-deoxycholate, 1% Nonidet-P40 and 1 mM EDTA) and LiCl Buffer (50 mM Tris-HCl, pH 8, 250 mM LiCl, 1 mM EDTA, 1% Nonidet-P40 and 0.5% Na- deoxycholate) and 1 ml TE (10 mM Tris-HCl, pH 8, 1 mM EDTA). Beads were transferred to fresh tubes, centrifuged and chromatin was eluted by incubating in 100 μl elution buffer (50 mM Tris-HCl, pH 8, 10 mM EDTA and 1% SDS) at 65°C with agitation. Chromatin was transferred to fresh tubes and incubated with 2 μl RNase A (Qiagen, 20 mg/ml) and 6 μl 5M NaCL (Active Motif) for 30 minutes at 37°C followed by 2 μl proteinase K (Active Motif, 0.2 mg/ml) at 65°C for 2 hours. DNA was then purified by phenol/chloroform extraction and resuspended in nuclease free water. Quantitative PCR was performed as described above. Primer sequences used were; FOXP3 CNS2, Forward: 5’-GGACATCACCTACCACATCC-3’ Reverse: 5’- ACCACGGAGGAAGAGAAGAG-3’; β-Actin, Forward: 5’-TCCCCTCCTTTTGCGAAAA-3’ Reverse: 5’- CTCCCTCCTCCTCTTCCTCAA -3’

### Statistical analyses

Statistical measures, including mean values, standard deviations, Student’s t-tests, Mantel–Cox tests, Mann–Whitney tests and one-way ANOVA tests, were performed using Graphpad Prism software and R. Definitions of n = values are stated in each figure legend. Where appropriate, unless otherwise stated, graphs display mean ± standard deviation.

## Acknowledgments

**General:** We would like to express our deep appreciation to Anne Hocking, Karen Cerosaletti, Jessica Hamerman and Daniel Campbell for helpful discussion. B10.Rag2^-/-^ mice were a kind gift from Dr. Brian Kelsall. We would like to acknowledge Tina Polintan for editorial assistance.

## Author contributions

BK, AA, RJX, PSL, VHG and TBS designed studies. AA, KGM, KJF, TBS, LJ, AFS and BK conducted experiments. AA, KJF, YZ and BK analyzed data. NSG, TD, YZ, DEL, IJM and ZSR provided reagents. AA and BK wrote the manuscript.

## Data and materials availability

RNAseq libraries generated in this study have been made available at the Gene Expression Omnibus (GEO) accession number: GSE141933

## References

1. E. Zigmond et al., Ly6C hi monocytes in the inflamed colon give rise to proinflammatory effector cells and migratory antigen-presenting cells. Immunity. 37, 1076–1090 (2012).

2. E. Zigmond, S. Jung, Intestinal macrophages: well educated exceptions from the rule. Trends Immunol. 34, 162–168 (2013).

3. S. Z. Josefowicz, L.-F. Lu, A. Y. Rudensky, Regulatory T cells: mechanisms of differentiation and function. Annu. Rev. Immunol. 30, 531–564 (2012).

4. E. Cretney, A. Kallies, S. L. Nutt, Differentiation and function of Foxp3(+) effector regulatory T cells. Trends Immunol. 34, 74–80 (2013).

5. J. J. O’shea, W. E. Paul, Mechanisms Underlying Lineage Commitment and Plasticity of Helper CD4+ T Cells. Science. 327, 1098–1102 (2010).

6. E. Batlle, J. Massagué, Transforming Growth Factor-β Signaling in Immunity and Cancer. Immunity. 50, 924–940 (2019).

7. B. Khor et al., The kinase DYRK1A reciprocally regulates the differentiation of Th17 and regulatory T cells. Elife. 4 (2015), doi:10.7554/eLife.05920.

8. T. B. Sundberg et al., Small-molecule screening identifies inhibition of salt-inducible kinases as a therapeutic strategy to enhance immunoregulatory functions of dendritic cells. Proc Natl Acad Sci USA (2014), doi:10.1073/pnas.1412308111.

9. M. Chen et al., Systemic Toxicity Reported for CDK8/19 Inhibitors CCT251921 and MSC2530818 Is Not Due to Target Inhibition. Cells. 8, 1413 (2019).

10. R. C. Conaway, J. W. Conaway, Function and regulation of the Mediator complex. Curr Opin Genet Dev. 21, 225–230 (2011).

11. S. Sato et al., A set of consensus mammalian mediator subunits identified by multidimensional protein identification technology. Mol Cell. 14, 685–691 (2004).

12. M. Malumbres, Cyclin-dependent kinases. Genome Biol. 15, 122 (2014).

13. M. D. Galbraith et al., CDK8 Kinase Activity Promotes Glycolysis. CellReports. 21, 1495–1506 (2017).

14. J. Bancerek et al., CDK8 kinase phosphorylates transcription factor STAT1 to selectively regulate the interferon response. Immunity. 38, 250–262 (2013).

15. A. Lin et al., Casein kinase II is a negative regulator of c-Jun DNA binding and AP-1 activity. Cell. 70, 777–789 (1992).

16. C.-C. Huang et al., Calcineurin-mediated dephosphorylation of c-Jun Ser-243 is required for c-Jun protein stability and cell transformation. Oncogene. 27, 2422–2429 (2008).

17. N. Taira et al., DYRK2 priming phosphorylation of c-Jun and c-Myc modulates cell cycle progression in human cancer cells. J Clin Invest. 122, 859–872 (2012).

18. M. Akamatsu et al., Conversion of antigen-specific effector/memory T cells into Foxp3- expressing Treg cells by inhibition of CDK8/19. Sci Immunol. 4, eaaw2707 (2019).

19. C. J. Fryer, J. B. White, K. A. Jones, Mastermind recruits CycC:CDK8 to phosphorylate the Notch ICD and coordinate activation with turnover. Mol Cell. 16, 509–520 (2004).

20. J. Shi et al., Scalable synthesis of cortistatin A and related structures. J Am Chem Soc. 133, 8014–8027 (2011).

21. H. E. Pelish et al., Mediator kinase inhibition further activates super-enhancer-associated genes in AML. Nature. 526, 273–276 (2015).

22. L. Johannessen et al., Small-molecule studies identify CDK8 as a regulator of IL-10 in myeloid cells. Nat. Chem. Biol. 13, 1102–1108 (2017).

23. A. Witalisz-Siepracka et al., NK Cell-Specific CDK8 Deletion Enhances Antitumor Responses. Cancer Immunol Res. 6, 458–466 (2018).

24. E. M. Putz et al., CDK8-mediated STAT1-S727 phosphorylation restrains NK cell cytotoxicity and tumor surveillance. CellReports. 4, 437–444 (2013).

25. Z. Guo, G. Wang, Y. Lv, Y. Y. Wan, J. Zheng, Inhibition of Cdk8/Cdk19 Activity Promotes Treg Cell Differentiation and Suppresses Autoimmune Diseases. Front Immunol. 10, 775–10 (2019).

26. J. Martinez-Fabregas et al., CDK8 Fine-Tunes IL-6 Transcriptional Activities by Limiting STAT3 Resident Time at the Gene Loci. CellReports. 33, 108545 (2020).

27. J. L. Coombes et al., A functionally specialized population of mucosal CD103+ DCs induces Foxp3+ regulatory T cells via a TGF-beta and retinoic acid-dependent mechanism. J Exp Med. 204, 1757–1764 (2007).

28. D. Mucida et al., Reciprocal TH17 and regulatory T cell differentiation mediated by retinoic acid. Science. 317, 256–260 (2007).

29. C. M. Sun et al., Small intestine lamina propria dendritic cells promote de novo generation of Foxp3 T reg cells via retinoic acid. J Exp Med. 204, 1775–1785 (2007).

30. S. Haxhinasto, D. Mathis, C. Benoist, The AKT-mTOR axis regulates de novo differentiation of CD4+Foxp3+ cells. J Exp Med. 205, 565–574 (2008).

31. J. A. Hill et al., Retinoic acid enhances Foxp3 induction indirectly by relieving inhibition from CD4+CD44hi Cells. Immunity. 29, 758–770 (2008).

32. S. Sauer et al., T cell receptor signaling controls Foxp3 expression via PI3K, Akt, and mTOR. Proc Natl Acad Sci USA. 105, 7797–7802 (2008).

33. J. A. Hall et al., Essential role for retinoic acid in the promotion of CD4(+) T cell effector responses via retinoic acid receptor alpha. Immunity. 34, 435–447 (2011).

34. A. E. Herman, G. J. Freeman, D. Mathis, C. Benoist, CD4+CD25+ T regulatory cells dependent on ICOS promote regulation of effector cells in the prediabetic lesion. J Exp Med. 199, 1479–1489 (2004).

35. K. V. Tarbell, S. Yamazaki, K. Olson, P. Toy, R. M. Steinman, CD25+ CD4+ T cells, expanded with dendritic cells presenting a single autoantigenic peptide, suppress autoimmune diabetes. J Exp Med. 199, 1467–1477 (2004).

36. F. Powrie, M. W. Leach, S. Mauze, L. B. Caddle, R. L. Coffman, Phenotypically distinct subsets of CD4+ T cells induce or protect from chronic intestinal inflammation in C. B-17 scid mice. Int Immunol. 5, 1461–1471 (1993).

37. V. Valatas et al., Host-dependent control of early regulatory and effector T-cell differentiation underlies the genetic susceptibility of RAG2-deficient mouse strains to transfer colitis. Mucosal immunology. 6, 601–611 (2013).

38. P. M. Smith et al., The microbial metabolites, short-chain fatty acids, regulate colonic Treg cell homeostasis. Science. 341, 569–573 (2013).

39. P. Kovarik et al., Specificity of signaling by STAT1 depends on SH2 and C-terminal domains that regulate Ser727 phosphorylation, differentially affecting specific target gene expression. EMBO J. 20, 91–100 (2001).

40. Y. Shen et al., Essential role of STAT3 in postnatal survival and growth revealed by mice lacking STAT3 serine 727 phosphorylation. Mol Cell Biol. 24, 407–419 (2004).

41. L. Varinou et al., Phosphorylation of the Stat1 transactivation domain is required for full-fledged IFN-gamma-dependent innate immunity. Immunity. 19, 793–802 (2003).

42. Z. Wen, Z. Zhong, J. E. Darnell, Maximal activation of transcription by Stat1 and Stat3 requires both tyrosine and serine phosphorylation. Cell. 82, 241–250 (1995).

43. M. Afkarian et al., T-bet is a STAT1-induced regulator of IL-12R expression in naïve CD4+ T cells. Nature Publishing Group. 3, 549–557 (2002).

44. A. A. Lighvani et al., T-bet is rapidly induced by interferon-gamma in lymphoid and myeloid cells. Proc Natl Acad Sci USA. 98, 15137–15142 (2001).

45. X. O. Yang et al., STAT3 regulates cytokine-mediated generation of inflammatory helper T cells. J Biol Chem. 282, 9358–9363 (2007).

46. Y. Zheng et al., Genome-wide analysis of Foxp3 target genes in developing and mature regulatory T cells. Nature. 445, 936–940 (2007).

47. A. Marson et al., Foxp3 occupancy and regulation of key target genes during T-cell stimulation. Nature. 445, 931–935 (2007).

48. W. Fu et al., A multiply redundant genetic switch “locks in” the transcriptional signature of regulatory T cells. Nat Immunol. 13, 972–980 (2012).

49. S. Hori, T. Nomura, S. Sakaguchi, Control of regulatory T cell development by the transcription factor Foxp3. Science. 299, 1057–1061 (2003).

50. W. Zheng, R. A. Flavell, The transcription factor GATA-3 is necessary and sufficient for Th2 cytokine gene expression in CD4 T cells. Cell. 89, 587–596 (1997).

51. E. A. Wohlfert et al., GATA3 controls Foxp3⁺ regulatory T cell fate during inflammation in mice. J Clin Invest. 121, 4503–4515 (2011).

52. G. Wei et al., Genome-wide analyses of transcription factor GATA3-mediated gene regulation in distinct T cell types. Immunity. 35, 299–311 (2011).

53. D. Rudra et al., Transcription factor Foxp3 and its protein partners form a complex regulatory network. Nat Immunol. 13, 1010–1019 (2012).

54. T. C. Fang et al., Notch directly regulates Gata3 expression during T helper 2 cell differentiation. Immunity. 27, 100–110 (2007).

55. C. Mota et al., Delta-like 1-mediated Notch signaling enhances the in vitro conversion of human memory CD4 T cells into FOXP3-expressing regulatory T cells. The Journal of Immunology. 193, 5854–5862 (2014).

56. N. Joller et al., Treg Cells Expressing the Coinhibitory Molecule TIGIT Selectively Inhibit Proinflammatory Th1 and Th17 Cell Responses. Immunity. 40, 569–581 (2014).

57. I. Yevshin, R. Sharipov, S. Kolmykov, Y. Kondrakhin, F. Kolpakov, GTRD: a database on gene transcription regulation-2019 update. Nucleic Acids Res. 47, D100–D105 (2019).

58. A. Subramanian et al., Gene set enrichment analysis: a knowledge-based approach for interpreting genome-wide expression profiles. Proc Natl Acad Sci USA. 102, 15545–15550 (2005).

59. V. K. Mootha et al., PGC-1alpha-responsive genes involved in oxidative phosphorylation are coordinately downregulated in human diabetes. Nat Genet. 34, 267–273 (2003).

60. M. Kitagawa, Notch signalling in the nucleus: roles of Mastermind-like (MAML) transcriptional coactivators. J Biochem. 159, 287–294 (2016).

61. Y. Wang, M. A. Su, Y. Y. Wan, An essential role of the transcription factor GATA-3 for the function of regulatory T cells. Immunity. 35, 337–348 (2011).

62. P.-Y. Mantel et al., GATA3-driven Th2 responses inhibit TGF-beta1-induced FOXP3 expression and the formation of regulatory T cells. PLoS Biol. 5, e329 (2007).

63. J. Wei et al., Antagonistic nature of T helper 1/2 developmental programs in opposing peripheral induction of Foxp3+ regulatory T cells. Proc Natl Acad Sci USA. 104, 18169–18174 (2007).

64. S. Hadjur et al., IL4 blockade of inducible regulatory T cell differentiation: the role of Th2 cells, Gata3 and PU.1. Immunol. Lett. 122, 37–43 (2009).

65. W. Ouyang et al., Inhibition of Th1 development mediated by GATA-3 through an IL-4- independent mechanism. Immunity. 9, 745–755 (1998).

66. R. Yagi et al., The Transcription Factor GATA3 Actively Represses RUNX3 Protein- Regulated Production of Interferon-&gamma. Immunity. 32, 507–517 (2010).

67. J. P. van Hamburg et al., Enforced expression of GATA3 allows differentiation of IL-17- producing cells, but constrains Th17-mediated pathology. Eur J Immunol. 38, 2573–2586 (2008).

68. D. F. Fiorentino, M. W. Bond, T. R. Mosmann, Two types of mouse T helper cell. IV. Th2 clones secrete a factor that inhibits cytokine production by Th1 clones. J Exp Med. 170, 2081–2095 (1989).

69. I. Menzl, A. Witalisz-Siepracka, V. Sexl, CDK8-Novel Therapeutic Opportunities. Pharmaceuticals. 12, 92–12 (2019).

70. R. Firestein et al., CDK8 is a colorectal cancer oncogene that regulates beta-catenin activity. Nature. 455, 547–551 (2008).

71. P. A. Clarke et al., Assessing the mechanism and therapeutic potential of modulators of the human Mediator complex-associated protein kinases. Elife. 5 (2016), doi:10.7554/eLife.20722.

72. J. Shi et al., Stereodivergent synthesis of 17-alpha and 17-beta-alpharyl steroids: application and biological evaluation of D-ring cortistatin analogues. Angew. Chem. Int. Ed. Engl. 48, 4328–4331 (2009).

73. D. J. Cousins, T. H. Lee, D. Z. Staynov, Cytokine Coexpression During Human Th1/Th2 Cell Differentiation: Direct Evidence for Coordinated Expression of Th2 Cytokines. J Immunol. 169, 2498–2506 (2002).

74. A. Chaudhry et al., Interleukin-10 signaling in regulatory T cells is required for suppression of Th17 cell-mediated inflammation. Immunity. 34, 566–578 (2011).

75. L. W. Collison, D. A. A. Vignali, In vitro Treg suppression assays. Methods Mol Biol. 707, 21–37 (2011).

76. Y. P. De Jong et al., Chronic murine colitis is dependent on the CD154/CD40 pathway and can be attenuated by anti-CD154 administration. Gastroenterology. 119, 715–723 (2000).

77. C. A. Schneider, W. S. Rasband, K. W. Eliceiri, NIH Image to ImageJ: 25 years of image analysis. Nat Methods. 9, 671–675 (2012).

78. A. Dobin et al., STAR: ultrafast universal RNA-seq aligner. Bioinformatics. 29, 15–21 (2013).

79. S. Anders, P. T. Pyl, W. Huber, HTSeq--a Python framework to work with high-throughput sequencing data. Bioinformatics. 31, 166–169 (2015).

80. H. Li et al., The Sequence Alignment/Map format and SAMtools. Bioinformatics. 25, 2078–2079 (2009).

81. H. Wickham et al., Welcome to the Tidyverse. JOSS. 4, 1686–6 (2019).

82. S. Durinck et al., BioMart and Bioconductor: a powerful link between biological databases and microarray data analysis. Bioinformatics. 21, 3439–3440 (2005).

83. D. J. McCarthy, Y. Chen, G. K. Smyth, Differential expression analysis of multifactor RNA-Seq experiments with respect to biological variation. Nucleic Acids Res. 40, 4288– 4297 (2012).

84. M. E. Ritchie et al., limma powers differential expression analyses for RNA-sequencing and microarray studies. 43, e47 (2015).

85. A. Liberzon et al., The Molecular Signatures Database (MSigDB) hallmark gene set collection. Cell Systems. 1, 417–425 (2015).

86. X. Xie et al., Systematic discovery of regulatory motifs in human promoters and 3’ UTRs by comparison of several mammals. Nature. 434, 338–345 (2005).

87. G. Seumois et al., Epigenomic analysis of primary human T cells reveals enhancers associated with TH2 memory cell differentiation and asthma susceptibility. Nat Immunol. 15, 777–788 (2014).

