## Supplemental Tables for "The CDK8 inhibitor DCA promotes a tolerogenic chemical immunophenotype in CD4^+^ T cells via a novel CDK8-GATA3-FOXP3 pathway"

**Supplementary Tables**

**Table S1.**

| | |
|---|---|
| Collection(s): | TFT:GTRD |
| # overlaps shown: | 100 |
| # genesets in collections: | 348 |
| # genes in comparison (n): | 556 |
| # genes in universe (N): | 40071 |

| Gene Set Name | # Genes in Gene Set (K) | # Genes in Overlap (k) | k/K | p-value | FDR q-value |
|---|---|---|---|---|---|
| MAML1_TARGET_GENES | 309 | 29 | 0.0939 | 8.34E-16 | 2.90E-13 |
| TFEB_TARGET_GENES | 1380 | 58 | 0.042 | 9.55E-14 | 1.66E-11 |
| HMG20B_TARGET_GENES | 1906 | 66 | 0.0346 | 1.07E-11 | 1.24E-09 |
| SKIL_TARGET_GENES | 1930 | 65 | 0.0337 | 5.03E-11 | 3.93E-09 |
| ZNF766_TARGET_GENES | 1191 | 48 | 0.0403 | 6.17E-11 | 3.93E-09 |
| ZNF597_TARGET_GENES | 875 | 40 | 0.0457 | 6.78E-11 | 3.93E-09 |
| ZNF768_TARGET_GENES | 1337 | 51 | 0.0381 | 1.06E-10 | 5.25E-09 |
| ZFP91_TARGET_GENES | 1436 | 53 | 0.0369 | 1.45E-10 | 6.32E-09 |
| E2F5_TARGET_GENES | 1267 | 46 | 0.0363 | 4.23E-09 | 1.41E-07 |
| NAB2_TARGET_GENES | 1491 | 51 | 0.0342 | 4.25E-09 | 1.41E-07 |
| ZNF423_TARGET_GENES | 1054 | 41 | 0.0389 | 4.47E-09 | 1.41E-07 |
| ZSCAN30_TARGET_GENES | 1738 | 56 | 0.0322 | 5.83E-09 | 1.69E-07 |
| GCM2_TARGET_GENES | 1974 | 60 | 0.0304 | 1.34E-08 | 3.59E-07 |
| HES2_TARGET_GENES | 1414 | 48 | 0.0339 | 1.57E-08 | 3.91E-07 |
| ZNF175_TARGET_GENES | 835 | 34 | 0.0407 | 3.25E-08 | 7.55E-07 |
| PCGF1_TARGET_GENES | 539 | 26 | 0.0482 | 5.42E-08 | 1.18E-06 |
| FOXN3_TARGET_GENES | 1490 | 47 | 0.0315 | 1.93E-07 | 3.96E-06 |
| SFMBT1_TARGET_GENES | 1643 | 50 | 0.0304 | 2.27E-07 | 4.39E-06 |
| GREB1_TARGET_GENES | 934 | 33 | 0.0353 | 1.25E-06 | 2.30E-05 |
| TERF1_TARGET_GENES | 293 | 16 | 0.0546 | 4.17E-06 | 7.26E-05 |
| ZNF592_TARGET_GENES | 1877 | 51 | 0.0272 | 4.57E-06 | 7.57E-05 |
| MAFG_TARGET_GENES | 1887 | 51 | 0.027 | 5.29E-06 | 8.36E-05 |
| RYBP_TARGET_GENES | 1843 | 50 | 0.0271 | 5.93E-06 | 8.97E-05 |
| POU2AF1_TARGET_GENES | 917 | 31 | 0.0338 | 6.38E-06 | 9.25E-05 |
| UBN1_TARGET_GENES | 1768 | 48 | 0.0271 | 9.00E-06 | 1.25E-04 |
| HHEX_TARGET_GENES | 1028 | 33 | 0.0321 | 9.36E-06 | 1.25E-04 |
| ZNF92_TARGET_GENES | 1434 | 41 | 0.0286 | 1.27E-05 | 1.64E-04 |
| ZNF512_TARGET_GENES | 438 | 19 | 0.0434 | 1.50E-05 | 1.86E-04 |
| ZNF350_TARGET_GENES | 1462 | 41 | 0.028 | 1.99E-05 | 2.38E-04 |
| RBM34_TARGET_GENES | 1023 | 32 | 0.0313 | 2.12E-05 | 2.46E-04 |
| CHAF1B_TARGET_GENES | 978 | 31 | 0.0317 | 2.22E-05 | 2.46E-04 |
| DACH1_TARGET_GENES | 979 | 31 | 0.0317 | 2.26E-05 | 2.46E-04 |
| ZNF589_TARGET_GENES | 683 | 24 | 0.0351 | 3.84E-05 | 4.04E-04 |

| | | | | | |
|---|---|---|---|---|---|
| PRKDC_TARGET_GENES | 873 | 28 | 0.0321 | 4.48E-05 | 4.59E-04 |
| FOXD2_TARGET_GENES | 1240 | 35 | 0.0282 | 7.09E-05 | 7.05E-04 |
| SALL4_TARGET_GENES | 1824 | 46 | 0.0252 | 8.16E-05 | 7.89E-04 |
| FOXJ2_TARGET_GENES | 587 | 21 | 0.0358 | 8.99E-05 | 8.45E-04 |
| ZFHX3_TARGET_GENES | 1843 | 45 | 0.0244 | 2.00E-04 | 1.83E-03 |
| HOXB6_TARGET_GENES | 628 | 21 | 0.0334 | 2.26E-04 | 2.01E-03 |
| LHX9_TARGET_GENES | 1552 | 39 | 0.0251 | 3.05E-04 | 2.60E-03 |
| SUPT16H_TARGET_GENES | 1935 | 46 | 0.0238 | 3.07E-04 | 2.60E-03 |
| KMT2D_TARGET_GENES | 793 | 24 | 0.0303 | 3.59E-04 | 2.98E-03 |
| ZNF197_TARGET_GENES | 652 | 21 | 0.0322 | 3.70E-04 | 3.00E-03 |
| NFKBIA_TARGET_GENES | 1313 | 34 | 0.0259 | 4.35E-04 | 3.44E-03 |
| PAX7_TARGET_GENES | 710 | 22 | 0.031 | 4.56E-04 | 3.45E-03 |
| SNIP1_TARGET_GENES | 759 | 23 | 0.0303 | 4.65E-04 | 3.45E-03 |
| ZIM3_TARGET_GENES | 1161 | 31 | 0.0267 | 4.66E-04 | 3.45E-03 |
| NCOA2_TARGET_GENES | 527 | 18 | 0.0342 | 4.84E-04 | 3.47E-03 |
| FEV_TARGET_GENES | 1645 | 40 | 0.0243 | 4.88E-04 | 3.47E-03 |
| ZNF322_TARGET_GENES | 934 | 26 | 0.0278 | 7.22E-04 | 5.03E-03 |
| E2F2_TARGET_GENES | 1464 | 36 | 0.0246 | 7.59E-04 | 5.18E-03 |
| ZNF563_TARGET_GENES | 643 | 20 | 0.0311 | 7.77E-04 | 5.20E-03 |
| TAZ_TARGET_GENES | 474 | 16 | 0.0338 | 1.10E-03 | 7.24E-03 |
| ZNF410_TARGET_GENES | 619 | 19 | 0.0307 | 1.21E-03 | 7.82E-03 |
| ZNF22_TARGET_GENES | 1186 | 30 | 0.0253 | 1.33E-03 | 8.43E-03 |
| PAX3_TARGET_GENES | 1799 | 41 | 0.0228 | 1.43E-03 | 8.91E-03 |
| ZNF843_TARGET_GENES | 881 | 24 | 0.0272 | 1.52E-03 | 9.26E-03 |
| ZNF274_TARGET_GENES | 1094 | 28 | 0.0256 | 1.61E-03 | 9.56E-03 |
| IGLV5_37_TARGET_GENES | 635 | 19 | 0.0299 | 1.62E-03 | 9.56E-03 |
| F10_TARGET_GENES | 275 | 11 | 0.04 | 1.75E-03 | 1.01E-02 |
| ZNF213_TARGET_GENES | 592 | 18 | 0.0304 | 1.79E-03 | 1.02E-02 |
| ZNF184_TARGET_GENES | 1486 | 35 | 0.0236 | 1.84E-03 | 1.03E-02 |
| ASH1L_TARGET_GENES | 1436 | 34 | 0.0237 | 1.96E-03 | 1.08E-02 |
| ZSCAN2_TARGET_GENES | 903 | 24 | 0.0266 | 2.09E-03 | 1.12E-02 |
| ZNF618_TARGET_GENES | 1442 | 34 | 0.0236 | 2.09E-03 | 1.12E-02 |
| MEF2D_TARGET_GENES | 651 | 19 | 0.0292 | 2.14E-03 | 1.12E-02 |
| ZBTB12_TARGET_GENES | 1279 | 31 | 0.0242 | 2.15E-03 | 1.12E-02 |
| ZNF596_TARGET_GENES | 654 | 19 | 0.0291 | 2.25E-03 | 1.15E-02 |
| SMN1_SMN2_TARGET_GENES | 917 | 24 | 0.0262 | 2.54E-03 | 1.28E-02 |
| ZNF513_TARGET_GENES | 766 | 21 | 0.0274 | 2.70E-03 | 1.34E-02 |
| PDCD6_AHRR_TARGET_GENES | 1359 | 32 | 0.0235 | 2.85E-03 | 1.40E-02 |
| ZNF418_TARGET_GENES | 43 | 4 | 0.093 | 2.95E-03 | 1.43E-02 |
| ARID5B_TARGET_GENES | 930 | 24 | 0.0258 | 3.03E-03 | 1.45E-02 |
| TEAD2_TARGET_GENES | 1483 | 34 | 0.0229 | 3.25E-03 | 1.53E-02 |
| ZNF282_TARGET_GENES | 1048 | 26 | 0.0248 | 3.51E-03 | 1.63E-02 |

| | | | | | |
|---|---|---|---|---|---|
| SNRNP70_TARGET_GENES | 1002 | 25 | 0.025 | 3.87E-03 | 1.77E-02 |
| PGM3_TARGET_GENES | 264 | 10 | 0.0379 | 4.09E-03 | 1.85E-02 |
| ZNF2_TARGET_GENES | 1177 | 28 | 0.0238 | 4.44E-03 | 1.98E-02 |
| LCORL_TARGET_GENES | 451 | 14 | 0.031 | 4.62E-03 | 2.03E-02 |
| ZSCAN31_TARGET_GENES | 916 | 23 | 0.0251 | 5.07E-03 | 2.21E-02 |
| ZNF30_TARGET_GENES | 1532 | 34 | 0.0222 | 5.30E-03 | 2.25E-02 |
| ZNF223_TARGET_GENES | 319 | 11 | 0.0345 | 5.37E-03 | 2.25E-02 |
| ZNF10_TARGET_GENES | 814 | 21 | 0.0258 | 5.37E-03 | 2.25E-02 |
| SETD7_TARGET_GENES | 980 | 24 | 0.0245 | 5.74E-03 | 2.38E-02 |
| PHF21A_TARGET_GENES | 281 | 10 | 0.0356 | 6.27E-03 | 2.57E-02 |
| SIX1_TARGET_GENES | 376 | 12 | 0.0319 | 6.80E-03 | 2.75E-02 |
| ZNF391_TARGET_GENES | 944 | 23 | 0.0244 | 7.17E-03 | 2.87E-02 |
| ZNF7_TARGET_GENES | 895 | 22 | 0.0246 | 7.65E-03 | 3.01E-02 |
| PAX8_TARGET_GENES | 88 | 5 | 0.0568 | 7.70E-03 | 3.01E-02 |
| CBX7_TARGET_GENES | 954 | 23 | 0.0241 | 8.08E-03 | 3.12E-02 |
| KLF7_TARGET_GENES | 914 | 22 | 0.0241 | 9.61E-03 | 3.68E-02 |
| ID1_TARGET_GENES | 1255 | 28 | 0.0223 | 1.01E-02 | 3.83E-02 |
| ZFP3_TARGET_GENES | 354 | 11 | 0.0311 | 1.12E-02 | 4.20E-02 |
| ZNF512B_TARGET_GENES | 218 | 8 | 0.0367 | 1.16E-02 | 4.30E-02 |
| ZNF558_TARGET_GENES | 310 | 10 | 0.0323 | 1.20E-02 | 4.33E-02 |
| TERF2_TARGET_GENES | 136 | 6 | 0.0441 | 1.20E-02 | 4.33E-02 |
| ATF5_TARGET_GENES | 456 | 13 | 0.0285 | 1.21E-02 | 4.33E-02 |
| ZNF502_TARGET_GENES | 35 | 3 | 0.0857 | 1.25E-02 | 4.44E-02 |
| HMGA1_TARGET_GENES | 1222 | 27 | 0.0221 | 1.28E-02 | 4.51E-02 |
| ZNF610_TARGET_GENES | 886 | 21 | 0.0237 | 1.31E-02 | 4.58E-02 |

4

**Table S2. Reagents Used**

| REAGENT | SOURCE | IDENTIFIER |
|---|---|---|
| Antibodies | | |
| Rabbit anti-Lamin B1 (clone D4Q4Z), unconjugated | Cell Signaling | Cat# 12586, RRID:AB_2650517 |
| Rabbit anti-Notch1 ICD (clone D3B8), unconjugated | Cell Signaling | Cat# 4147, RRID:AB_2153348) |
| Mouse Anti-b-actin, clone (AC-15), unconjugated | Sigma Aldrich | Cat# A1978, RRID:AB_476692 |
| Rabbit anti-c-Jun, (clone 60A8), unconjugated | Cell Signaling | Cat# 9165, RRID:AB_2130165 |
| Rat anti-CD25 (clone PC61.5), Alexa 488 | eBioscience | Cat# 14-0251-86, RRID:AB_467177 |
| Syrian Hamster anti-CD28 (clone 37.51), unconjugated | BioXCell | Cat# BE0015-1, RRID: AB_1107624 |
| Armenian Hamster anti-CD3 (clone 145-2C11), unconjugated | BioXCell | Cat# BE0001-1, RRID:AB_1107634 |
| Rat anti-CD4 (clone RM4-5), FITC, PerCP | Biolegend | Cat# 100505, RRID:AB_312709 |
| Rat anti-CD45RB (clone C363.16A), APC | eBioscience | Cat# 12-0455-82, RRID:AB_465681 |
| Rat anti-CD62L (clone MEL-14), PE | eBioscience | Cat# 12-0621-82, RRID:AB_465721 |
| Rat anti-Foxp3 (clone FJK-16s), PE | eBioscience | Cat# 14-5773-82, RRID:AB_467576 |
| Rat anti-IFNγ (clone XMG1.2), unconjugated | BioXCell | Cat# BE0055, RRID:AB_1107694 |
| Rat anti-IFNγ (clone XMG1.2), APC | eBioscience | Cat# 17-7311-82, RRID:AB_469504 |
| Rat anti-IL-12 (clone C17.8), unconjugated | BioXCell | Cat# BE0051, RRID:AB_1107698 |
| Rat anti-IL-17A (clone eBio17B7), APC | eBioscience | Cat# 17-7177-81, RRID:AB_763580 |
| Rat anti-IL4 (clone 11B11), unconjugated | BioXCell | Cat# BE0045, RRID:AB_1107707 |
| Rat anti-IL4 (BVD6-24G2), PE-Cyanine7 | eBioscience | Cat#25-7042-41, RRID:AB_1659723 |
| Rabbit polyclonal anti-Mouse, HRP | Cell Signaling | Cat# 7074, RRID:AB_2099233 |
| Rabbit poly clonal anti-phospho-c-Jun (pS243), unconjugated | Cell Signaling | Cell Signaling Technology Cat# 2994, RRID:AB_823470 |

| | | |
|---|---|---|
| Rabbit polyclonal Anti-phospho-c-Jun (pS63) unconjugated | Cell Signaling | Cat# 9261, RRID:AB_2130162 |
| Rabbit anti-phospho-S6 (clone D57.2.2E) Alexa647 | Cell Signaling | Cat# 4851, RRID:AB_10695457 |
| Rabbit anti-phospho-S6 kinase (clone Poly9205), unconjugated | Cell Signaling | Cat# 9205, RRID:AB_330944 |
| Rabbit anti-phospho-Smad2 (clone 138D4), unconjugated | Cell Signaling | Cat# 3108, RRID:AB_490941 |
| Rabbit Anti-phospho-Smad3 (clone C25A9), unconjugated | Cell Signaling | Cat# 9520, RRID:AB_2193207 |
| Rabbit Anti-phospho-Stat1 (S727) (clone D3B7) Alexa647, unconjugated | Cell Signaling | Cat# 8826, RRID:AB_2773718) |
| Mouse anti-phospho-Stat1 (Y701) (clone 4a), Alexa488, unconjugated | BD Biosciences | Cat# 612596, RRID:AB_399879 |
| Rabbit Anti-phospho-Stat3 (S727) (clone D8C2Z), unconjugated | Cell Signaling | Cat# 94994, RRID:AB_2800239 |
| Rabbit Anti-phospho-Stat3 (Y705) (clone D3A7), PE | Cell Signaling | Cat# 9145, RRID:AB_2491009 |
| Rabbit polyclonal anti-phospho-Stat5b (S731), unconjugated | Abcam | Cat# ab52211, RRID:AB_2196931 |
| Rat anti-RORγt (clone AFKJS-9), APC | eBioscience | Cat# 17-6988-82, RRID:AB_10609207 |
| Mouse anti-S6 kinase (clone H9), unconjugated | Santa Cruz | Cat# sc-8418, RRID:AB_628094 |
| Rabbit Anti-Smad2 (clone D43B4), unconjugated | Cell Signaling | Cat# 5339, RRID:AB_10626777 |
| Rabbit Anti-Smad3 (clone C67H9), unconjugated | Cell Signaling | Cat# 9523, RRID:AB_2193182 |
| Rabbit Anti-Stat1 (clone D1K9Y), Alexa647, unconjugated | Cell Signaling | Cat# 14994, RRID:AB_2737027 |
| Rabbit Anti-Stat3 (clone 79D7), Alexa647, unconjugated | Cell Signaling | Cat# 4904, RRID:AB_331269 |
| Rabbit Anti-Stat5 (clone D2O6Y), unconjugated | Cell Signaling | Cat# 94205, RRID:AB_2737403 |
| Mouse anti-T-bet (clone 4B10), PE | Biolegend | Cat# 644809, RRID:AB_2028583 |
| Mouse Anti-CD127 (clone eBioRDR5), APC | eBioscience | Cat# 17-1278-42, RRID:AB_1659670 |

| | | |
|---|---|---|
| Mouse anti-CD28 (clone 28.2), unconjugated | Biolegend | Cat# 302902, RRID:AB_314304 |
| Mouse anti-CD3 (clone UCHT), unconjugated | BioXCell | Cat# BE0231, RRID:AB_2687713 |
| Mouse anti-CD4 (clone RPA-T4), PerCP | eBioscience | Cat# 45-0049-42, RRID:AB_1518744 |
| Mouse anti-CD45RA (clone L48), FITC | BD Biosciences | Cat# 347723, RRID:AB_400343 |
| Mouse anti-CD45RO (clone UCHL1), PE | eBioscience | Cat# 12-0457-42, RRID:AB_1272079 |
| Mouse anti-Foxp3 (clone 236A/E7), PE | eBioscience | Cat# 12-4777-42, RRID:AB_1944444 |
| Rabbit Anti-GATA3 (clone D13C9), unconjugated | Cell Signaling | Cat# 5852, RRID:AB_10835690 |
| Mouse anti-IFNγ (clone B27), unconjugated | Biolegend | Cat# 506502, RRID:AB_315435 |
| Mouse anti-IL-12/IL-23 p40 (clone C8.6), unconjugated | Biolegend | Cat# 508807, RRID:AB_2810643 |
| Rat anti-Il-4 (clone MP4-25D2), PE | BD Biosciences | Cat# 554485, RRID:AB_395424 |
| Rat anti-Il-4 (clone MP4-25D2), unconjugated | Biolegend | Cat# 500802, RRID:AB_315121 |
| Mouse anti-NGFR (clone ME20.4), FITC | Biolegend | Cat# 345103, RRID:AB_1937226 |
| Rabbit Normal rabbit IgG (clone 2729), unconjugated | Cell Signaling | Cat# 2729, RRID:AB_1031062 |
| **Bacterial and Virus Strains** | | |
| Lenti-pLKO-NGFR-T2A-GATA3 | This paper | N/A |
| Lenti-pLKO-NGFR-T2A-c-JUN | This paper | N/A |
| Lenti-pLKO-NGFR-T2A-JUNB | This paper | N/A |
| Lenti-pLKO-NGFR-T2A-JUND | This paper | N/A |
| Lenti-pLKO-NGFR | Chance John lucky, This paper | N/A |
| **Biological Sample** | | |
| Human peripheral mononuclear cells | BRI Repository | https://www.benaroyaresearch.org/what-is-bri/scientists-and-laboratories/core-labs/translational-core-laboratory |
| **Chemical Compounds, Peptides, and Recombinant Proteins** | | |
| All-trans retinoic acid | Sigma Aldrich | R2625; CAS: 302-79-4, |
| Rapamycin | Cayman Chemical | 13346; CAS:53123-88-9 |
| Didehydro-cortistatin a | P. Baran, Shi J et al., 2011 | N/A |
| T5224 | Cayman Chemical | 22904; CAS:530141-72-1 |
| HG 9-91-01 | T. Sundberg, Sundberg TB et al., 2014 | N/A |
| Harmine | Cayman Chemical | 10010324; CAS: 442-51-3 |

| | | |
|---|---|---|
| Human TGFβ | PeproTech | 100-21; Accession number: P01137 |
| Human Il-4 | PeproTech | 200-04; Accession number: P05112 |
| Human Il-2 | PeproTech | 200-02; Accession number: P60568 |
| Murine Il-2 | PeproTech | 212-12; Accession number: P04351 |
| Murine Il-4 | PeproTech | 214-14; Accession number: P07750 |
| Murine Il-12 | PeproTech | 210-12; Accession numbers: P43431, P43432 |
| Murine Il-6 | PeproTech | 216-16; Accession number: P08505 |
| Murine Il-1β | PeproTech | 211-11b; Accession number: P10749 |
| Commercial Assay, Kit | | |
| EasySep Human Naïve CD4+ T Cell Isolation Kit II | Stem Cell | Cat# 17555 |
| EasySep Mouse CD4+ T Cell Isolation Kit | Stem Cell | Cat# 19852 |
| CD4+ CD62L+ T Cell Isolation Kit, mouse | Miltenyi Biotec | Cat# 130-093-227 |
| Gibson Assembly ultra-kit | Codex | Cat# GA-1200 |
| RNeasy Kit | Qiagen | Cat# 74106 |
| iScript DNA synthesis Kit | BioRad | Cat# 1708890 |
| Nuclei EZ prep Kit | Sigma Aldrich | Cat# NUC101-1KT |
| SMART-Seq v4 Ultra low inout RNA Kit | Takara | Cat# 634894 |
| NexteraXT DNA sample preparation Kit | Illumina | Cat# FC 131-1096 |
| SBS kit | Illumina | Cat# FC-402-4022 |
| Large Scale Data set | | |
| RNASeq Libraries | GEO | GSE141933 |
| Cell line | | |
| Human: 293T | David Rawlings, Certo MT et al., 2011 | N/A |
| Organisms/Strains | | |
| Murine: Balb/c, | The Jackson Laboratory | Stock No: 000651 RRID:IMSR_JAX:000651 |
| Murine: *Foxp3*$^{IRES\text{-}GFP}$ | The Jackson Laboratory | Stock No: 006772 RRID:IMSR_JAX:006772 |
| Murine: *CD45.1*$^{+/+}$ | The Jackson Laboratory | Stock No: 002014RRID:IMSR_JAX:002014 |
| Murine: C57Bl/6 | The Jackson Laboratory651 | Stock No: 000664RRID:IMSR_JAX:000664 |
| C57Bl/10-*Rag2*$^{-/-}$ | Brian Kelsall, Valatas V et al | N/A |
| Murine: *Stat1*$^{Ser727Ala}$ | Ziaur Rahman, Varinou L et al., 2003 | N/A |

| | | |
|---|---|---|
| Murine: *Stat3*$^{Ser727Ala}$ | David Levy, Shen Y et al., 2004 | N/A |
| Murine: NOD-*BDC2.5.Foxp3*$^{IRES-GFP}$ | JDRF Transgenic Core | N/A |
| Murine: NOD-*scid* | JDRF Transgenic Core | N/A |
| Oligonucleotides | | |
| Primer FOXP3 CNS2 Forward: 5'-GGACATCACCTACCACATCC-3' | IDT, This paper | N/A |
| Primer FOXP3 CNS2 Reverse: : 5'-ACCACGGAGGAAGAGAAGAG-3' | IDT, This paper | N/A |
| Primer b-Actin Forward: 5'-TCCCCTCCTTTTGCGAAAA-3' | IDT, This paper | N/A |
| Primer b-Actin Reverse: 5'-CTCCCTCCTCCTCTTCCTCAA -3' | IDT, This paper | N/A |
| gRNA 1 CDK8: CUCAUGCUGAUAGGAAG | Synthego, This paper | N/A |
| gRNA 2 CDK8: UGUUUCUGUCUCAUGCUGAU | Synthego, This paper | N/A |
| gRNA 3 CDK8: UCUGUCUCAUGCUGAUAGGA | Synthego, This paper | N/A |
| For primers used for gene expression analysis, see Supplementary file 1 | IDT, This paper | N/A |
| For primers for plasmid preparation, see Supplementary file 1 | IDT, This paper | N/A |
| Recombinant DNA reagent | | |
| pLKO-NGFR-T2A-GATA3 | This paper | N/A |
| pLKO-NGFR-T2A-c-JUN | This paper | N/A |
| pLKO-NGFR-T2A-JUNB | This paper | N/A |
| pLKO-NGFR-T2A-JUND | This paper | N/A |
| pLKO-NGFR | Chance John Lucky, This paper | N/A |
| pHAGE-GATA3 | Ng et al,. 2018 | Addgene Plasmid #116747 |
| JunD-HA neo | Wang CC et al., 2014 | Addgene Plasmid # 58515 |
| pMIEG3-c-Jun | Wang ZY et al., 2005 | Addgene Plasmid # 40348 |
| pMIEG3-JunB | Wang ZY et al., 2005 | Addgene Plasmid # 40349 |
| pMD2.G | David Rawlings, Certo MT et al., 2011 | N/A |
| psPax2 | David Rawlings, Certo MT et al., 2011 | N/A |
| Software and Algorithm | | |
| MSigDB v7.0 | Subramanian et al., 2005 | https://www.gsea-msigdb.org/gsea/msigdb |
| ImageJ | Schneider et al., 2012 | https://imagej.nih.gov/ij/ |
| Prism 7 | Graphpad | https://www.graphpad.com/scientific-software/prism/ |

| | | |
|---|---|---|
| Illumina real time analysis | Illumina HiSeq 2500 | https://www.illumina.com/systems/sequencing-platforms/hiseq-2500.htm |
| R v3.6.1 | R Core Team 2019 | https://www.r-project.org |
| limma v3.40.6 | Ritchie et al., 2015 | http://bioconductor.org/packages/release/bioc/html/limma.html |
| biomart v2.40.5 | Durinck et al., 2009 | https://bioconductor.org/packages/release/bioc/html/biomaRt.html |
| dplyr v1.0.2 | Wickham et al., 2019 | https://dplyr.tidyverse.org |
| STARv.2.4.2a | Dobin et al. 2013 | http://code.google.com/p/rna-star/ |
| edgeR v3.26.8 | McCarthy et al., 2012 | http://bioconductor.org/packages/release/bioc/html/edgeR.html |
| stringr v1.4.0 | Wickham et al., 2019 | https://dplyr.tidyverse.org |
| samtools v1.9 | Li et al., 2009 | http://samtools.sourceforge.net/ |
| Picard v1.128 | N/A | http://broadinstitute.github.io/picard/ |
| FastQC | N/A | https://www.bioinformatics.babraham.ac.uk/projects/fastqc/ |
| htseq v0.3.7 | Anders et al., 2014 | https://htseq.readthedocs.io/en/master/ |

### Table S3. Primers

| | | |
|---|---|---|
| | Primers used in cloning | |
| | Primers to Amplify Insert and Partial T2A sequence | |
| **Insert** | **Forward** | **Reverse** |
| **GATA3** | TCTCCTTACTTGTGGCGACGTGGAGGAGAACCCCGGCCCCATGGAGGTGACGGCGG | TGGCGGTGACCATGCT |
| **c-JUN** | TCTCCTTACTTGTGGCGACGTGGAGGAGAACCCCGGCCCCATGACTGCAAAGATGGAAACG | AAATGTTTGCAACTGCTGCGT |
| **JUNB** | CCTTACTTGTGGCGACGTGGAGGAGAACCCCGGCCCCATGTGCACGAAAATGGAACAGCC | TCAGAAGGCGTGTCCCT |
| **JUND** | TCTCCTTACTTGTGGCGACGTGGAGGAGAACCCCGGCCCCATGGAAACGCCCTTCTATGG | CGGGACCTGGTGCTG |
| | Primers to add Homology Arms | |
| **Insert** | **Forward** | **Reverse** |
| **GATA3** | TGTGGTTGTGGGCCTTGTGGCCTACATAGCCTTCAAGAGG | TTTGTAATCCAGAGGTTGATTATCGATAAGCTTGATCTAACCCATGGCGGTGACCATGCT |
| **c-JUN** | TGTGGTTGTGGGCCTTGTGGCCTACATAGCCTTCAAGAGGTCC | TTTGTAATCCAGAGGTTGATTATCGATAAGCTTGATAAATGTTTGCAACTGCTGCGT |
| **JUNB** | TGTGGTTGTGGGCCTTGTGGCCTACATAGCCTTCAAGAGGTCCGGATCC | AAATTTTGTAATCCAGAGGTTGATTATCGATAAGCTTGATTCAGAAGGCGTGTCCCT |
| **JUND** | TGTGGTTGTGGGCCTTGTGGCCTACATAGCCTTCAAGAGGTCCGGATCC | TTTGTAATCCAGAGGTTGATTATCGATAAGCTTGATTTAGTACGCCGGGACCTGGTGCTG |
| | Primers for Remaining T2A sequence | |
| **Insert** | **Forward** | **Reverse** |
| **GATA3** | TTCAAGAGGTCCGGATCCGGAGAGGGCAGGGGATCTCTCCTTACTTGTGGCGAC | TGGCGGTGACCATGCT |
| **c-JUN** | TTCAAGAGGTCCGGATCCGGAGAGGGCAGGGGATCTCTCCTTACTTGTGGCGAC | AAATGTTTGCAACTGCTGCGT |

| | | |
|---|---|---|
| **JUNB** | TTCAAGAGGTCCGGATCCGGAGAGGGCAGGG GATCTCTCCTTACTTGTGGCGAC | TCAGAAGGCGTGTCCCT |
| **JUND** | TTCAAGAGGTCCGGATCCGGAGAGGGCAGGG GATCTCTCCTTACTTGTGGCGAC | CGGGACCTGGTGCTG |
| | **Primers used in qRT-PCR** | |
| **Target** | **Forward** | **Reverse** |
| **Foxp3** | GGCCCTTCTCCAGGACAGA | GCTGATCATGGCTGGGTTG |
| **Ikzf2** | TCACAACTATCTCCAGAATGTCAGC | AGGCGGTACATGGTGACTCAT |
| **Ikzf4** | CGGCATCCGGCTACCCAACG | AGGTCACGGATTTCATCACCTGGC |
| **Il2ra** | CCACATTCAAAGCCCTCTCCTA | GTTTTCCCACACTTCATCTTGC |
| **Il2** | TTGTGCTCCTTGTCAACAGC | CTGGGGAGTTTCAGGTTCC |
| **CDK8** | GACTATCAGCGTTCCAATCCAC | TAGCTGAGTATCCCATGCTGC |
| **β-ACTIN** | CACCATTGGCAATGAGCGGTTC | AGGTCTTTGCGGATGTCCACGT |
| **RPS18** | TCATCCTCCGTGAGTTCTCCA | AGTTCCAGCACATTTTGCGAG |
| **GATA3** | ACCACAACCACACTCTGGAGGA | TCGGTTTCTGGTCTGGATGCCT |
| **Gata3** | TGCCGACAGCCTTCGCTTGG | CCAAGGCACGATCCAGCACAGA |

**Table S4. Conditions Used**

| Condition | Species | Stimulation (μg/ml) | Neutralizing antibodies (μg/ml) | Cytokines (ng/ml) | Compound (μM) |
|---|---|---|---|---|---|
| $T_{reg}^{hi}$ | Mouse | anti-CD3 (1)<br>anti-CD28 (1) | Anti-IL-12 (2)<br>Anti-IFNγ (2)<br>Anti-IL-4 (2) | TGFβ (10) | |
| $T_{reg}^{low}$ | Mouse | anti-CD3 (1)<br>anti-CD28 (1) | Anti-IL-12 (2)<br>Anti-IFNγ (2)<br>Anti-IL-4 (2) | TGFβ (2) | |
| $T_{reg}^{low+DCA}$ | Mouse | anti-CD3 (1)<br>anti-CD28 (1) | Anti-IL-12 (2)<br>Anti-IFNγ (2)<br>Anti-IL-4 (2) | TGFβ (2) | DCA (0.1) |
| $Th17^{hi}$ | Mouse | anti-CD3 (1-3)<br>anti-CD28 (1-3) | Anti-IL-12 (2)<br>Anti-IFNγ (2)<br>Anti-IL-4 (2) | TGFb (0.25-0)<br>IL-6 (20)<br>IL-1b (20) | |
| $Th17^{low}$ | Mouse | anti-CD3 (1-3)<br>anti-CD28 (1-3) | Anti-IL-12 (2)<br>Anti-IFNγ (2)<br>Anti-IL-4 (2) | TGFb (0.25-0)<br>IL-6 (5) | |
| $Th1^{hi}$ | Mouse | anti-CD3 (1)<br>anti-CD28 (1) | -<br>- | IL-12 (10) | |
| $Th1^{low}$ | Mouse | anti-CD3 (1)<br>anti-CD28 (1) | -<br>- | IL-12 (0.25) | |
| $Th2^{hi}$ | Mouse | anti-CD3 (1)<br>anti-CD28 (1) | Anti-IL-12 (2)<br>Anti-IFNγ (2) | Il-4 (100) | |
| $Th2^{low}$ | Mouse | anti-CD3 (1)<br>anti-CD28 (1) | Anti-IL-12 (2)<br>Anti-IFNγ (2) | Il-4 (10) | |
| $T_{reg}^{hi}$ | Human | anti-CD3 (1)<br>anti-CD28 (1) | Anti-IL-12 (2)<br>Anti-IFNγ (2)<br>Anti-IL-4 (2) | TGFβ (10) | |
| $T_{reg}^{low}$ | Human | anti-CD3 (1)<br>anti-CD28 (1) | Anti-IL-12 (2)<br>Anti-IFNγ (2)<br>Anti-IL-4 (2) | TGFβ (1) | |
| $T_{reg}^{low+DCA}$ | Human | anti-CD3 (1)<br>anti-CD28 (1) | Anti-IL-12 (2)<br>Anti-IFNγ (2)<br>Anti-IL-4 (2) | TGFβ (1) | DCA (0.1) |
| $Th2^{low}$ | Human | anti-CD3 (1)<br>anti-CD28 (1) | Anti-IL-12 (10)<br>Anti-IFNγ (10) | Il-4 (10) | |
