## Supplemental Figures for "The CDK8 inhibitor DCA promotes a tolerogenic chemical immunophenotype in CD4^+^ T cells via a novel CDK8-GATA3-FOXP3 pathway"

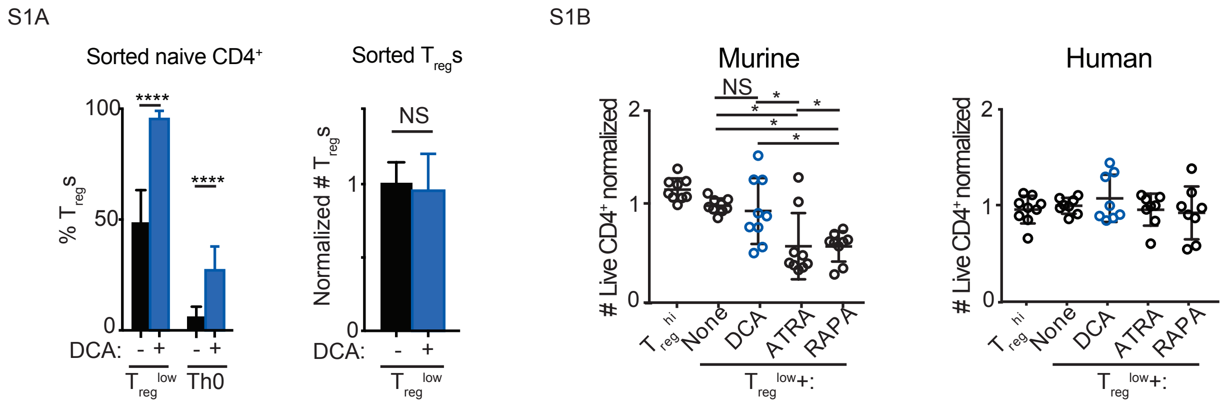


**Supplemental Fig. 1. DCA enhances differentiation, not proliferation, of murine $T_{reg}$s.** (A) Effect of DCA on $T_{reg}$ differentiation and proliferation in FACS-sorted murine naive $CD4^+$ T cells (left) and $T_{reg}$s (right) (≥4, x4 experiments). (B) Comparison of how DCA, ATRA and RAPA affect survival of murine (n = 9, x4 experiments) and human (n = 7-8, x3 experiments) $CD4^+$ T cells cultured in $T_{reg}^{low}$ conditions. Mann-Whitney (A-B), * P<0.05, ****P<0.0001.

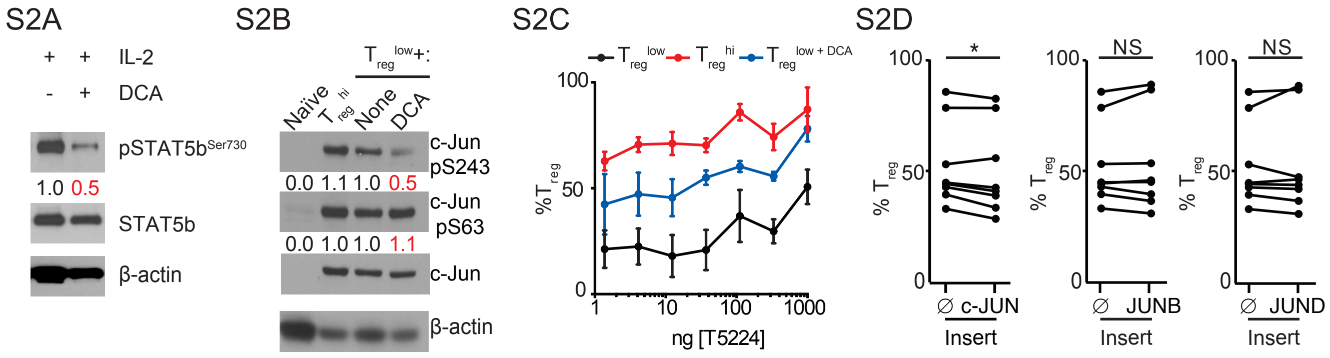


**Supplemental Fig. 2 DCA enhances $T_{reg}$ differentiation independent of regulating AP-1 transcription factor activity.** (A) Effect of DCA on IL-2-induced $STAT5b^{Ser730}$ phosphorylation in resting murine $CD4^+$ T cells (representative of 2 independent experiments). (B) Effect of DCA on c-Jun phosphorylation at Ser243 and Ser63 (representative of 3 independent experiments). (C) Effect of the c-Fos inhibitor T5224 on DCA's ability to enhance $T_{reg}$ differentiation in murine $CD4^+$ T cells cultured in $T_{reg}^{low}$ conditions. (n = 4, representative of 2 experiments). (D) Effect of overexpressing AP-1 family members using NGFR-T2A-tagged lentivirus and NGFR (∅) control on $T_{reg}$ differentiation in human $CD4^+$ T cells cultured for 5 days in $T_{reg}^{low}$ conditions (n = 8, x2 experiments). Wilcoxon matched pair analysis, * P<0.05.

S3A

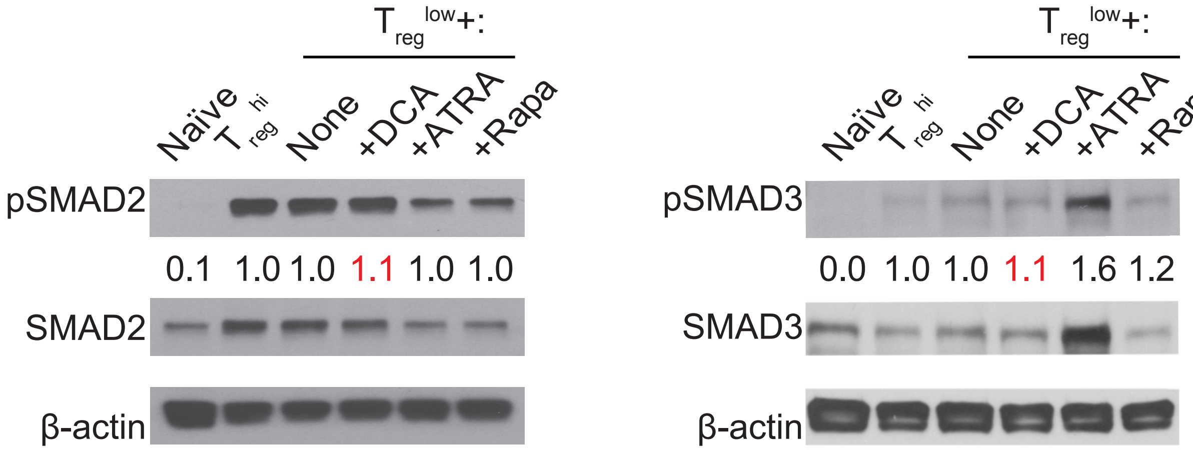


S3B

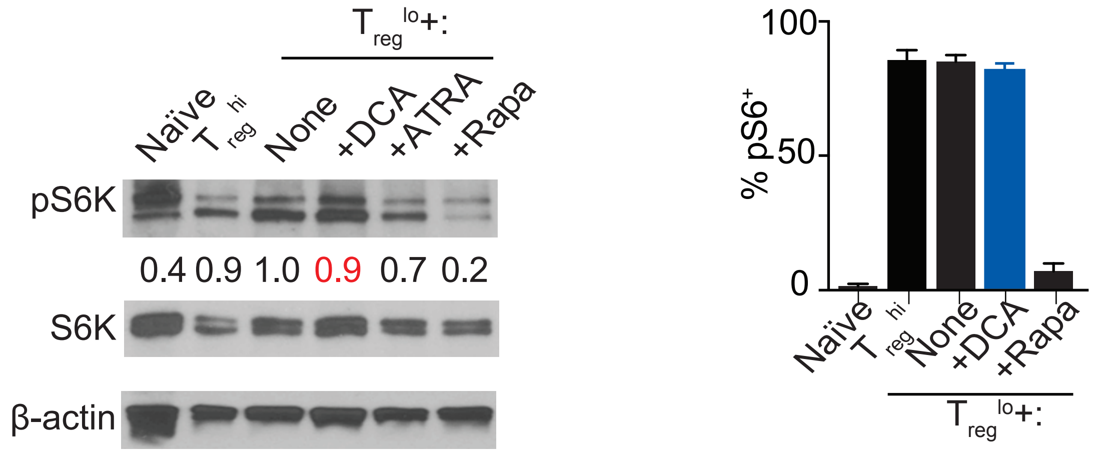


**Supplemental Fig. 3. CDK8 inhibition does not impact canonical pathways of $T_{reg}$ differentiation.** (A) Effect of DCA on phosphorylation of SMAD2 and SMAD3 in murine $CD4^{+}$ T cells stimulated in indicated conditions (≥2 independent experiments). (B) Effect of DCA on mTOR activity, as assessed by Western blot analysis of S6K phosphorylation (left) and flow cytometric analysis of S6 phosphorylation (right) in murine $CD4^{+}$ T cells stimulated in the indicated conditions. (A-B) Numbers in Western blot analyses show fraction of protein phosphorylated normalized to $T_{reg}^{low}$ conditions (representative of at least 2 experiments).

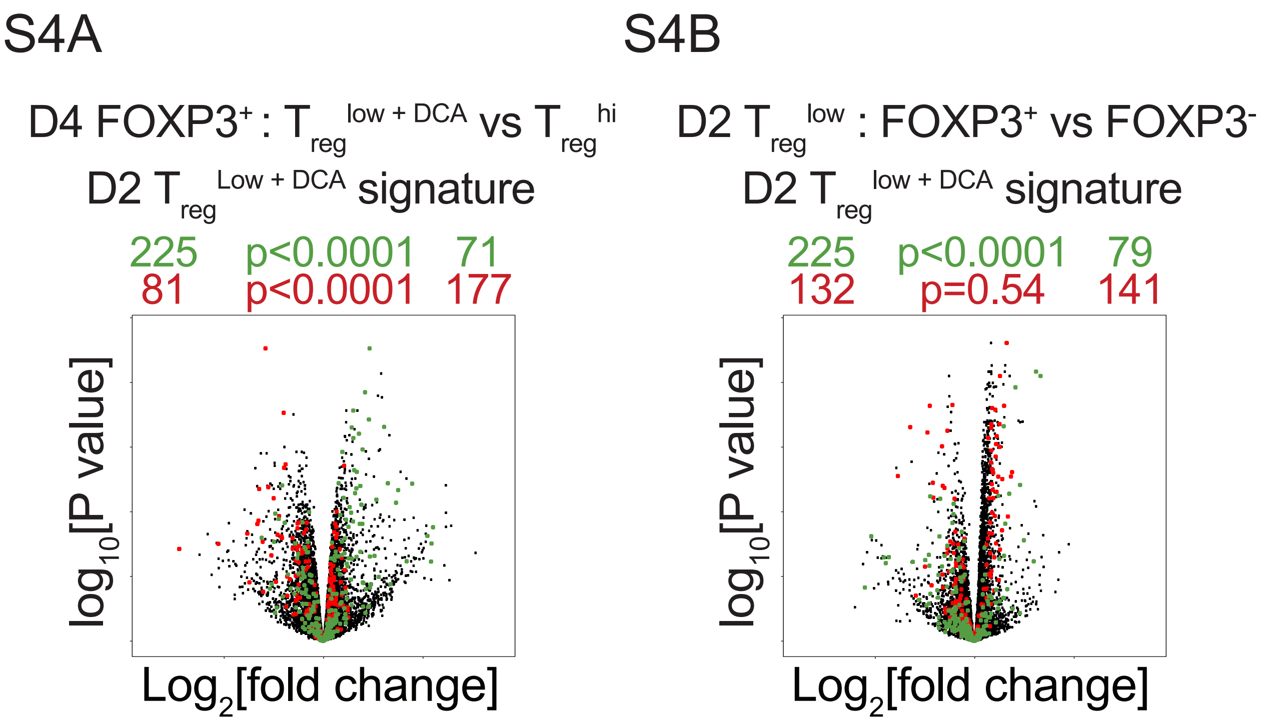


**Supplemental Fig. 4. Analysis of the DCA-associated transcriptional signature.** Volcano plots comparing gene expression in (A) sorted mature murine $T_{reg}$s generated in $T_{reg}^{low+DCA}$ versus $T_{reg}^{hi}$ conditions, and (B) murine CD4$^+$ T cells cultured for 2 days in $T_{reg}^{low}$ conditions, sorted FOXP3$^+$ versus FOXP3$^-$ cells. DCA signature genes, obtained by comparing sorted FOXP3$^+$ cells cultured for 2 days in $T_{reg}^{low+DCA}$ versus $T_{reg}^{low}$ conditions are highlighted in green (downregulated) and red (upregulated). Numbers on the right and left reflect genes that are up- and down-regulated in the indicated comparison respectively with $\chi 2$ test p values in the middle.
